# Two levels of selection of rhythmicity in gene expression: energy saving for rhythmic proteins and noise optimization for rhythmic transcripts

**DOI:** 10.1101/2021.04.15.439944

**Authors:** David Laloum, Marc Robinson-Rechavi

## Abstract

Many genes have nycthemeral rhythms of expression, *i.e.* a 24-hours periodic variation, at either mRNA or protein level or both, and, most rhythmic genes are tissue-specific. Here, we investigate and discuss the evolutionary origins of rhythms in gene expression. Our results suggest that thythmicity of protein expression has been favored by selection for low cost. Trends are consistent in bacteria, plants and animals, and are also supported in tissue-specific patterns in mouse. Cost cannot explain rhythm at the RNA level, and we suggest instead it allows to periodically and drastically reduce expression noise. Noise control had strongest support in mouse, with limited power in other species. Genes under stronger purifying selection are rhythmically expressed at the mRNA level, probably because they are noise sensitive genes. We also suggest that mRNA rhythmicity allows to switch between optimal precision and higher stochasticity. Higher precision allows to maximize the robustness of gene expression when the function is most needed, while higher stochasticity allows to maintain oscillations and to exhibit diverse molecular phenotypes, i.e. “blind anticipation” of cells. The ability to alternate between these two states, enabled by rhythmicity at the mRNA level, might be adaptive in fluctuating environments. Finally, the adaptive role of rhythmic expression is also supported by rhythmic genes being highly expressed yet tissue-specific genes. This provides a good evolutionary explanation for the observation that nycthemeral rhythms are often tissue-specific.

## Introduction

Living organisms have to adapt to complex and changing environments. In a given environment, some conditions vary over time in predictable ways. Physiological systems able to accommodate themselves to changing circumstances are expected to have a higher stability of survival and reproduction (Graham 1997). The strongest predictable change for most organisms is the light/dark cycle and the associated nycthemeral temperature variations. In an organism, “circadian rhythms” denotes entities characterized by an endogenous and entrainable oscillator clock which is able to persist in constant conditions (such as in constant darkness) and whose phases can be altered (reset or entrained). However, many physiological systems display non-autonomous nycthemeral rhythms, directly or indirectly controlled by the local clock or mainly by the environment itself, or by both (Hubbard et al. 2020, Yoo et al. 2004, Boothroyd et al. 2007, Nagoshi et al. 2004, Gerber et al. 2015). Such rhythms are found at all levels: molecular, cellular, organs, and behavioural, and several regulatory networks appear to play roles in the synchronization of these levels (Saini et al. 2011).

Here, we studied the evolutionary costs and benefits that shape the rhythmic nature of gene expression at the RNA and protein levels. For this, we analysed characteristics we presume to be part of the trade-off determining the rhythmic nature of gene expression between its advantages (cost economy over 24h, ribosomal non-occupancy) and disadvantages (costs of complexity due to precise temporal regulation). We call “rhythmic genes” all genes displaying a 24-hours periodic variation of their mRNA or protein level, or of both, constituting the nycthemeral transcriptome or proteome. Their rhythmic expression can be entrained directly by the internal clock but also directly or indirectly by external inputs, such as the light-dark cycle or food-intake (Hubbard et al. 2020, Yoo et al. 2004, Boothroyd et al. 2007, Nagoshi et al. 2004, Gerber et al. 2015). Hence we use the term “nycthemeral” to avoid confusion with the specific features of “circadian” rhythms, although these are included in the nycthemeral rhythms. Because the alternation of light and dark can be considered as a permanent signal for most life on earth (Policarpo et al. 2020), we consider that the entirety of nycthemeral biological rhythms are relevant as a phenotype under selection.

### Rhythmic gene expression: an adaptation in cycling environments

The endogenous nature of circadian rhythms is a strategy of anticipation. The ability to anticipate improves the adaptation of organisms to their fluctuating environment. Most rhythmic genes are tissue-specific (Zhang et al. 2014, Boyle et al. 2017, Korenčič et al. 2014), *i.e.* a given gene can be rhythmic in some tissues, and constantly or not expressed in others, which means that their rhythmic regulation is not a general property of the gene and is therefore expected to be advantageous only in those tissues in which they are found rhythmic. This argues that rhythmic regulation has costs, since it is not general. These costs are probably related to the complexity of regulation to maintain precise temporal organisation. Thus, cyclic biological systems are expected to have adaptive origins. Gene expression is costly for the cell in terms of energy and cellular materials usage. Wang et al. (2015) provided first results showing that in the liver of mice, abundant proteins that are required at one time are down-regulated at other times, apparently to economize on overall production. Wang et al. also reported that at each time-point, the total metabolic cost was ~4 fold higher for the set of cycling genes compared to the non-cycling genes set at both transcriptional and translational levels (Wang et al. 2015) – although the proteomic data used from mouse fibroblasts appear to have been underestimated and have since been corrected (Schwanhäusser et al. 2013). Furthermore, we have shown in our previous work that rhythmic genes are largely enriched in highly expressed genes and that the differences in rhythm detection obtained between highly and lowly expressed genes either reflect true biology or a lower signal to noise ratio in lowly expressed genes (Laloum & Robinson-Rechavi 2020). Here, we present results supporting the hypothesis that cyclic expression of highly expressed proteins were selected as part of a “low-cost” strategy to minimize the overall use of cellular energy. Thus, a first evolutionary advantage given by rhythmic biological processes would be an optimization of the overall cost (over a 24-hour period), compared to the costs generated over the same period by optimizing a constant level of proteins. Furthermore, we provide a first explanation for the tissue-specificity of rhythms in gene expression by showing that genes are more likely to be rhythmic in tissues where they are specifically highly expressed.

### Noise and cost optimization

Expression costs at the protein level are at least 160 times higher than at the RNA level (section 3 in Supporting information). Thus, it requires considerably more metabolic activity to produce a significant change in protein than in transcript levels. Furthermore, small increases in expression levels have been shown to incur energy costs large enough to be opposed by natural selection, at least in bacteria (Wagner 2005, Lynch & Marinov 2015). Consequently, protein synthesis costs are more constraining than RNA synthesis costs, especially in eukaryotes for which translation is the major contribution to making expression costs visible to selection (Lynch & Marinov 2015).

Many transcripts show nycthemeral fluctuations without rhythmicity of their protein abundances, even when measured in the same study. For instance in mouse liver (Mauvoisin et al. 2014) or in plants (Baerenfaller et al. 2012). Yet comparatively to protein synthesis costs, costs at RNA level are probably too small to provide a satisfactory evolutionary explanation of rhythms at mRNA level.

Thattai et al. (2001) and Hausser et al. (2019) propose that living systems have made tradeoffs between energy efficiency and noise reduction. Indeed, the control of noise (stability against fluctuations) plays a key role in the function of biological systems (*i.e.* in the robustness of gene expression) as supported by observations of a noise reduction during key periods such as in some developmental stages (Liu et al. 2020, Thattai & van Oudenaarden 2001). These considerations lead to predictions which we test here: i) a strategy to periodically decrease stochasticity for genes with rhythmically accumulated mRNAs; ii) a cost-saving strategy for genes whose protein expression is rhythmic; and iii) a combined strategy for genes rhythmic at both levels.

## Results

### Cyclicality of costliest genes

Expression costs at the protein level are at least 160 times higher than at the RNA level (section 3 in Supporting information). They are dominated by the cost of translation (Lynch & Marinov 2015, Wagner 2005). This is mainly due to the higher abundance of proteins relative to transcripts, by a factor of 1000 (section 3 in Supporting information). Thus, we estimated the expression cost (*C*_*p*_) of each gene by the formula (*1*) which takes into account the averaged amino-acid (AA) synthesis cost 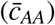 of the protein (Supp Table S2), protein length (*L*_*p*_), and protein abundance (*N*_*p*_). Other costs such as AA polymerization or protein decay are not expected to change our results, since longer or higher expressed proteins need more chain elongation activity and degradation (section 4 in Supporting information). We show in Supporting information (section 4) that the costs of maintaining proteins is higher for highly expressed than for lowly expressed proteins, even taking into account that highly expressed proteins have longer half-lives (also see Discussion). Thus, the cost provided by formula (*1*) is representative of gene expression costs and can be used for comparison between genes. To compare expression costs between rhythmic and non-rhythmic proteins, we calculated the average and the maximum protein expression level over time-points (see Methods equations (2) and (3)) (Figure 1d). The AA biosynthesis costs estimated in *E. coli* (Supplementary Table S2) were used as representative for all species since biosynthetic pathways are nearly universally conserved (Wagner 2005).

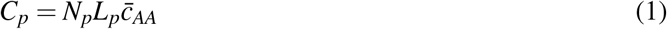

**Figure 1:**
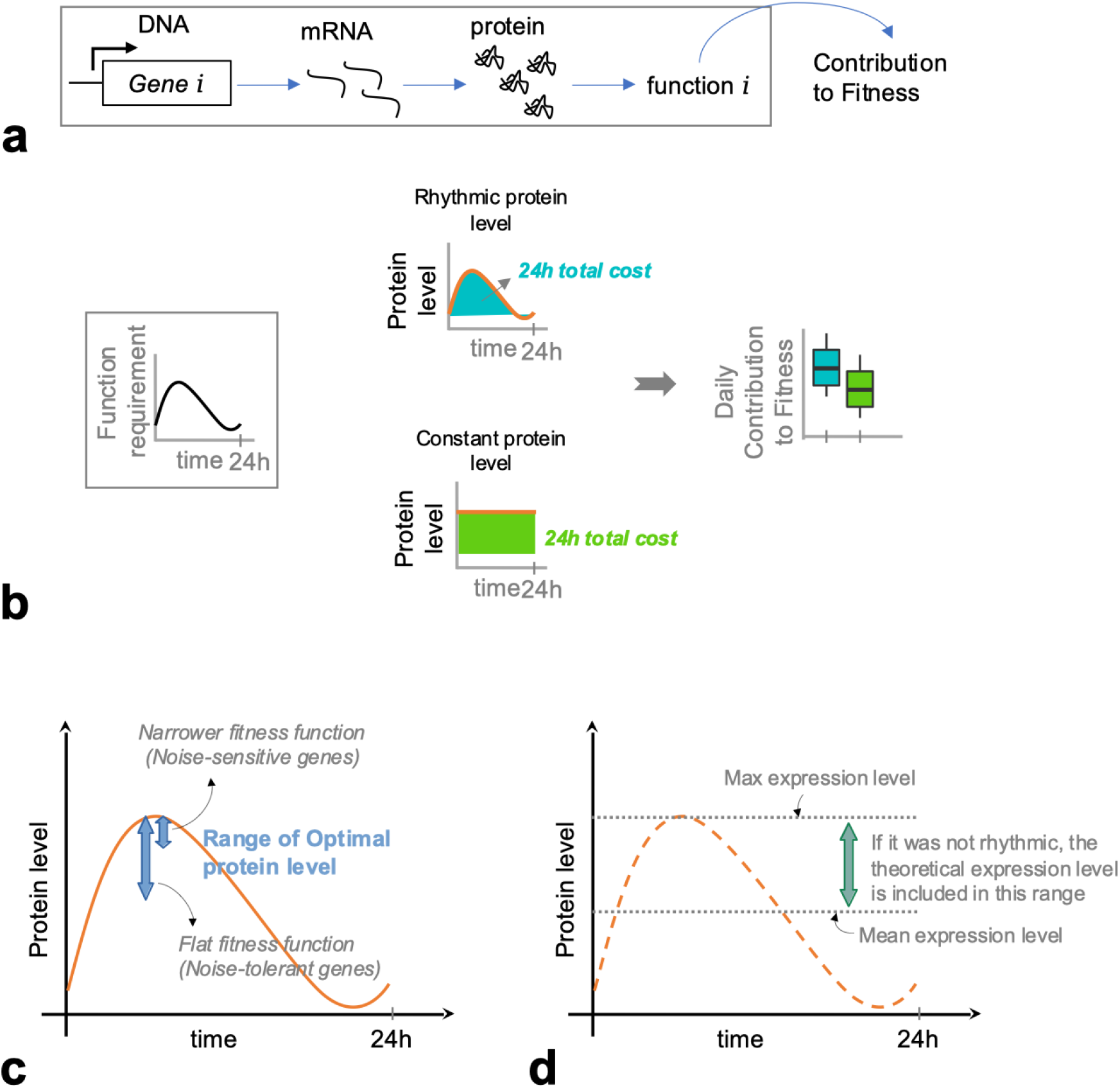
**a)** Gene expression contributes to organismal fitness. **b)** Rhythmic protein regulation presents a trade-off between the costs generated by integration into the rhythmic system (costs of complexity) and the advantages provided plus the costs saved over 24 hours. **c)** The range of high fitness protein levels depends on the sensitivity of the function to deviations from an optimal level. We use the terms “narrower” following Hausser et al. (2019). Noise sensitive genes have narrower fitness function, *i.e.* a small deviation from the optimum rapidly decreases the contribution to fitness. Precision is less important for genes with flat fitness functions. **d)** Mean or maximum expression level calculated from time-series datasets (see Methods). We assume that, in the absence of rhythmic regulation, the constant optimal level is included between the mean and the maximum expression level observed in rhythmic expression.

If the biological function of a protein is periodic, we expect the rhythmic regulation of its expression to be determined by a trade-off between the benefits of not producing proteins when they are not needed (costs saved) and the costs involved in making it rhythmic and temporally coordinated (costs of complexity) (Figure 1b). This leads to expect costlier genes to be more frequently rhythmic. First, we confirm that cycling genes are indeed enriched in highly expressed genes, as previously reported (Laloum & Robinson-Rechavi 2020, Wang et al. 2015). The higher cost of cycling genes (Figure 2a and Fig. S2) was especially due to their higher expression levels, observed both at the protein level (Figure 2d-e) and at the RNA level (Fig. S1). We find rhythmic proteins to be longer only in mouse and in cyanobacteria (Figure 2c).

**Figure 2:**
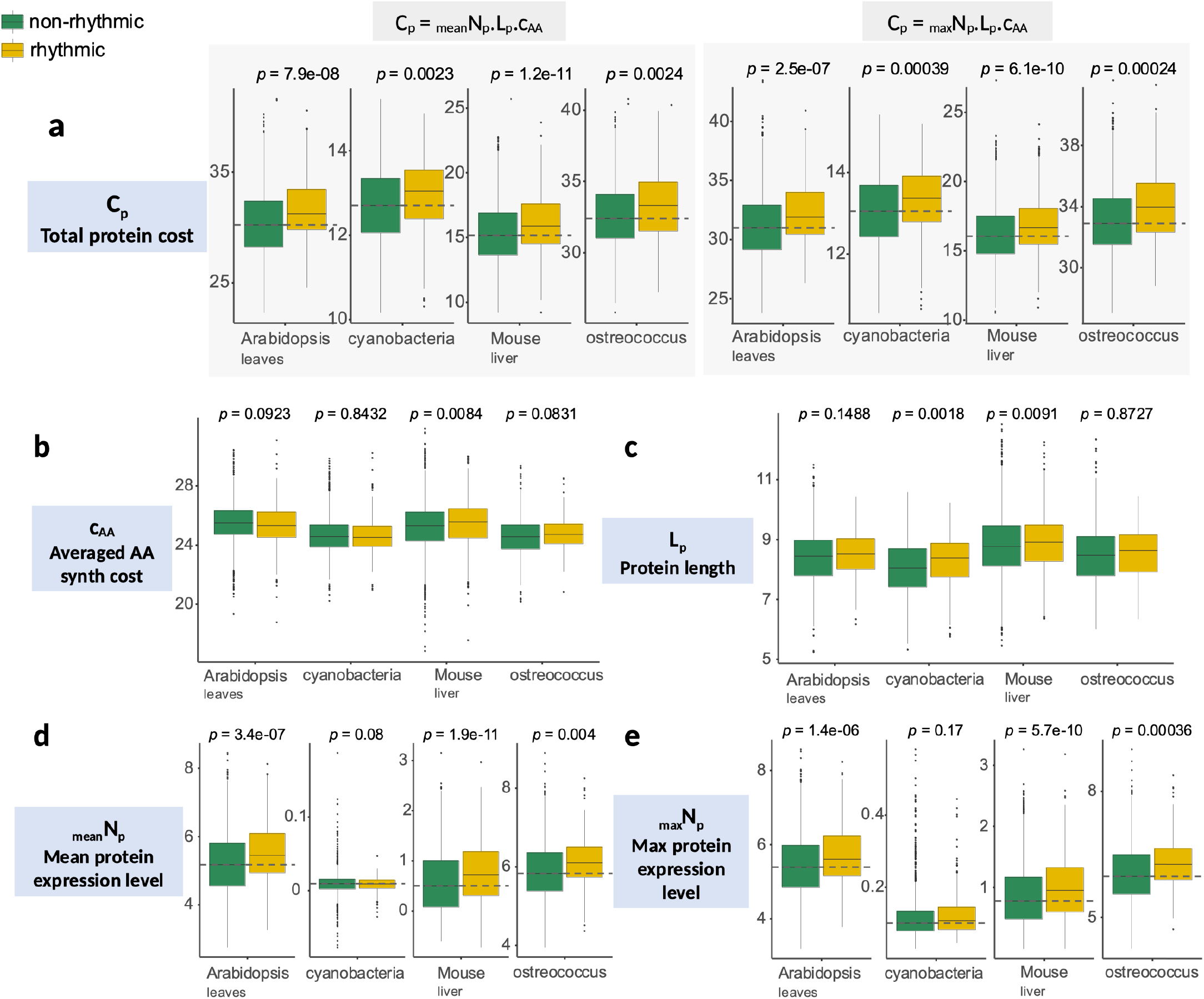
High level of expression is the main factor explaining the higher cost observed in rhythmic proteins. **a)** The total cost of rhythmic proteins is higher than those of other proteins. **b)** With the exception of mouse liver, rhythmic proteins do not contain more expensive amino-acids than other proteins. **c)** Rhythmic proteins can be longer in some species. **d-e)** Mean or maximum expression level calculated from time-series datasets: rhythmic proteins are highly expressed proteins. Boxplots are log scaled except for the averaged AA synthesis cost. The first 15% of proteins from *p*-values ranking obtained from the rhythm detection algorithms were classified as rhythmic.

However, conserved genes are known to be longer (Van Oss & Carvunis 2019, Neme & Tautz 2013) and are often more expressed (Wall et al. 2005, Drummond et al. 2005), therefore more expensive to produce, which could explain the rhythmic expression we observe for longer proteins. On the other hand, the higher cost for rhythmic proteins in *Synechococcus elongatus* appears only due to their length. Finally, cycling proteins seem to contain more expensive amino acids in mouse liver (Figure 2b).

Overall, results are consistent with the expectation that costlier genes are preferentially under rhythmic regulation. A more specific prediction from this hypothesis is that each gene should be found rhythmic specifically in tissues where a high expression of that gene is needed, *i.e.* where its function is costlier.

### Genes tend to be rhythmic in the tissues in which they are highly expressed

To test this hypothesis, we used a mouse circadian dataset with 11 tissues (transcriptomics from Zhang et al. (2014), Supplementary Table S1). For each gene we separated the tissues in two groups, those for which the gene was rhythmic (*p*-value≤0.01) and those for which it was not (*p*-value*>*0.5). Because of the difficulty of setting reliable thresholds for rhythmicity (Laloum & Robinson-Rechavi 2020), we ignored intermediate *p*-values. For each gene, we estimated the difference δ of expression levels between these two groups of tissues. As expected, genes tend to be more highly expressed in tissues where they are rhythmic (Student’s test, testing the hypothesis that the δ distribution mean is equal to 0: δ_*mean*_=0.0146, t=28.29, df=11683, *p*-value*<*2.2e^−16^, 95% CI [0.0136, 0.0156]). We also provide results obtained from other datasets in supplementary Table S3, although they must be taken with caution since only 2 to 4 tissues were available, and sometimes data were coming from different experiments. Of note, for proteomic data, the distributions of δ are bimodal (Fig. S3), separating rhythmic proteins into two groups, with low or high protein levels in the tissues in which they are rhythmic. A hypothesis is that for some tissue-specific proteins the rhythmic regulation is not tissue-specific, making them rhythmic also in tissues where they are lowly expressed. But the very small sample size does not allow us to test it, and we caution against any over-interpretation of this pattern before it can be confirmed.

### Rhythmic genes are tissue-specific

To clarify whether rhythmic genes tend to be tissue-specific highly expressed genes, we also analysed the relation between the number of tissues in which a gene is rhythmic and its tissue-specificity of expression τ (Yanai et al. 2004, Kryuchkova-Mostacci & Robinson-Rechavi 2016). For the mouse circadian dataset (transcriptomic) with 11 tissues, partial correlations show that genes whose rhythmicity is tissue-specific have tissue-specific expression (τ versus number of rhythmic tissues: Pearson’s correlation = −0.37, t=−5.3e+01, *p<*2.2e^−16^; see Supplementary Table S4 for Spearman’s correlation test). We provide results obtained from other datasets in supplementary Table S4 although they should be taken with caution for the baboon dataset for reasons discussed in our previous work (Laloum & Robinson-Rechavi 2020). The partial correlation seems to be stronger at the transcriptional level, although data was available in much less tissues at the protein level (only mouse forebrain, cartilage, liver, and tendon). Moreover, for tissue-specific genes (τ *>*0.5), the signal of rhythmicity correlates with expression level over tissues (Fig. S4a). While Spearman’s correlation is clearly skewed towards negative correlations, *i.e.* lower *p*-values thus stronger signal of rhythmicity in the tissue where genes are more expressed, Pearson’s correlation also has a smaller peak of positive correlations (Fig. S4a), suggesting a subset of genes which are less rhythmic in the tissues where they are most expressed. We show that tissue-specific genes which are mostly rhythmic in tissues where they are highly expressed are under stronger selective constraint than those which are rhythmic in tissues where they are lowly expressed (Fig. S4b).Thus, rhythmic expression of this second set of genes might be under weaker constraints. The dominant signal overall is that in a given tissue, rhythmically expressed genes tend to be those which are tissue-specifically expressed in this tissue.

### Lower cell-to-cell variability of genes with rhythmic transcripts

Increasing transcription for a fixed amount of protein can decrease the noise in final protein levels (Hausser et al. 2019), thus, genes with rhythmic mRNAs can have lower noise at their mRNA level peak. We predict that genes with rhythmic RNA have noise-sensitive functions (Fig. 1c). To test this, we compared the noise distribution between rhythmic and non-rhythmic genes. We used *F*\* of Barroso et al. (2018) to control for the correlation between expression mean (*μ*) and variance (σ^2^). It was the most efficient compared with other methods (section 6.4 in Supporting information and Supplementary Table S7). To evaluate expression noise, we used single-cell RNA data from *Mus musculus* liver, lung, limb muscle, heart, and aorta (Consortium 2018*a*) and *Arabidopsis thaliana* roots (Efroni et al. 2015) (Supplementary Table S1). In the absence of other data, we have to assume that this noise calculated at the transcriptional level is representative of noise at the protein level, or in any case functionally relevant. Then, we assessed rhythmicity based on time-series datasets: RNA and proteins from Arabidopsis leaves and mouse liver from the data used above; and RNA only in mouse lung, kidney, muscle, heart, and aorta (Supplementary Table S1). Results are given in a simplified way in Table 1 for the mouse; full results, which include Arabidopsis, are provided in supplementary Table S5. In most cases, we found lower noise for genes with rhythmic mRNA (Table 1a). Genes with both rhythmic proteins and mRNA had lower noise (Table 1c), although differences weren’t significant. Our results show that noise is globally reduced for genes with rhythmic regulation at the transcriptional level. Since rhythmic genes are not all in the same phase (Fig. S9a in Supporting information), we expect this result obtained for a given time-point (noise estimation based on a single time-point scRNA dataset) to be general to all time-points (section 6.3 in Supporting information). Assuming that genes with low noise have noise-sensitive functions (and thus noise is tightly controlled), these results suggest that rhythmic genes have their noise periodically and drastically reduced through periodic high accumulation of their mRNAs. In Arabidopsis, the single-cell data used are from the root, while transcriptomic time-series data used to detect rhythmicity are from the leaves, which limits the interpretation. Despite this limitation, we found no evidence of lower noise for genes that are rhythmic at the protein level (Table 1b and 1e, and Supplementary Table S5), and trends towards lower noise in almost all cases for genes with rhythmic mRNAs (Table 1a, 1c, and 1d).

**Table 1:**
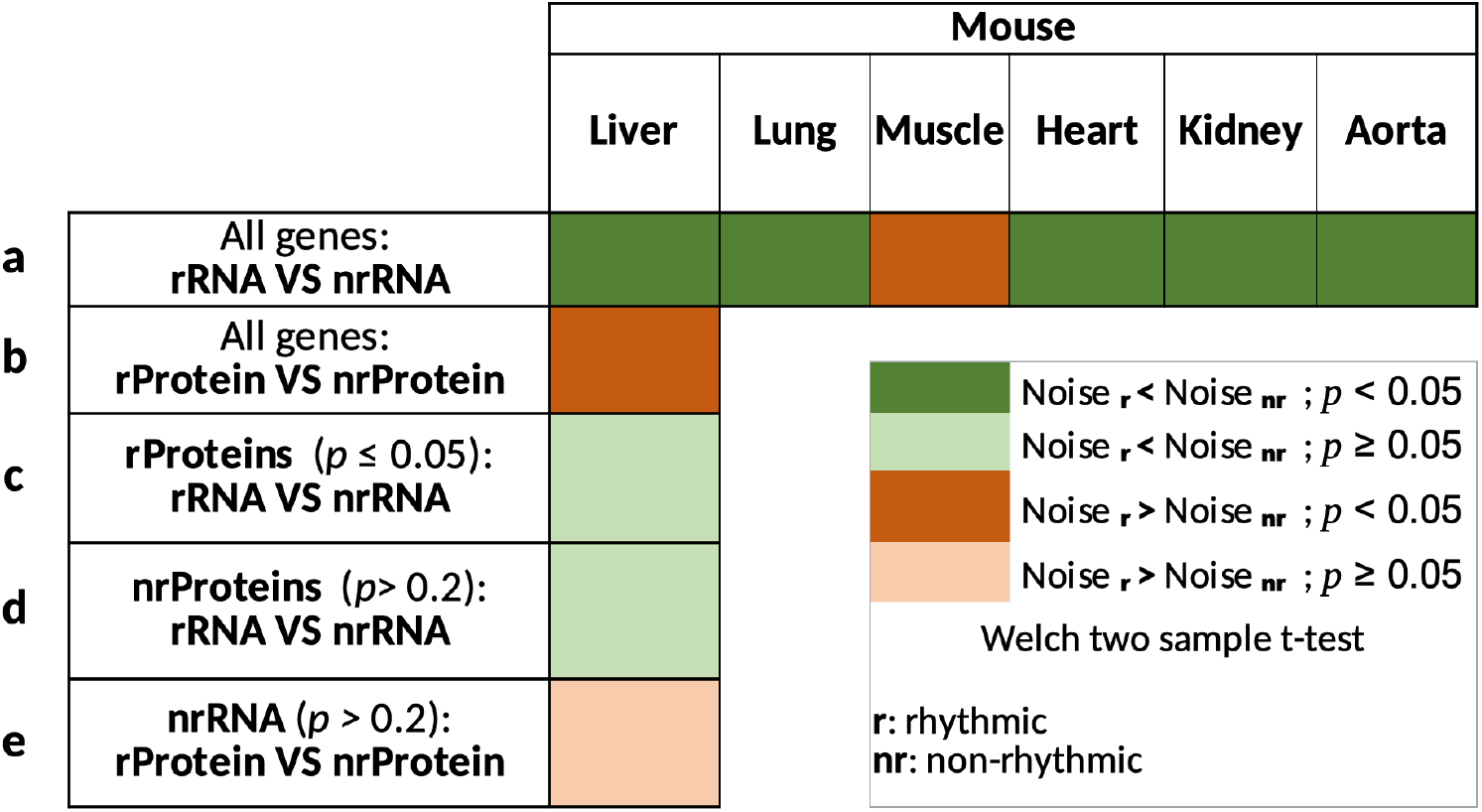
Simplified table showing the results of the Welch two sample t-test testing the hypothesis that the noise is equal between rhythmic versus non-rhythmic transcripts (**a**), or proteins (**b**), or between rhythmic versus non-rhythmic transcripts among rhythmic proteins (**c**) or among non-rhythmic proteins (**d**), and between rhythmic versus non-rhythmic proteins among genes with constant transcripts (**e**). *F*\* is an estimation of the noise based on Barroso et al. method (Barroso et al. 2018). Complete results are provided in Supplementary Table S5.

### Genes with rhythmic transcripts are under stronger selection

Finally, we compared protein evolutionary conservation (estimated by the dN/dS ratio) between rhythmic and non-rhythmic genes. Results are given in a simplified way in Table 2; full results are provided in supplementary Table S6. In all cases, in plants, vertebrates, and insects, we found that genes with rhythmic mRNA levels were significantly more conserved (triangles in Table 2a). We obtained similar results after controlling for gene expression bias (background boxes in Table 2a), since higher expressed genes are known to be more conserved genes. Interestingly, we didn’t obtain such clear results for rhythmic proteins (Table 2b and 2d). In Arabidopsis, genes rhythmic at both levels were less conserved than genes rhythmic at only one level (Table 2b and 2c), and results were unclear for mouse. These results suggest that rhythmic expression at the transcriptional level plays an important role for genes under strong purifying selection, consistent with the results for tissue-specific rhythmic genes.

**Table 2:**
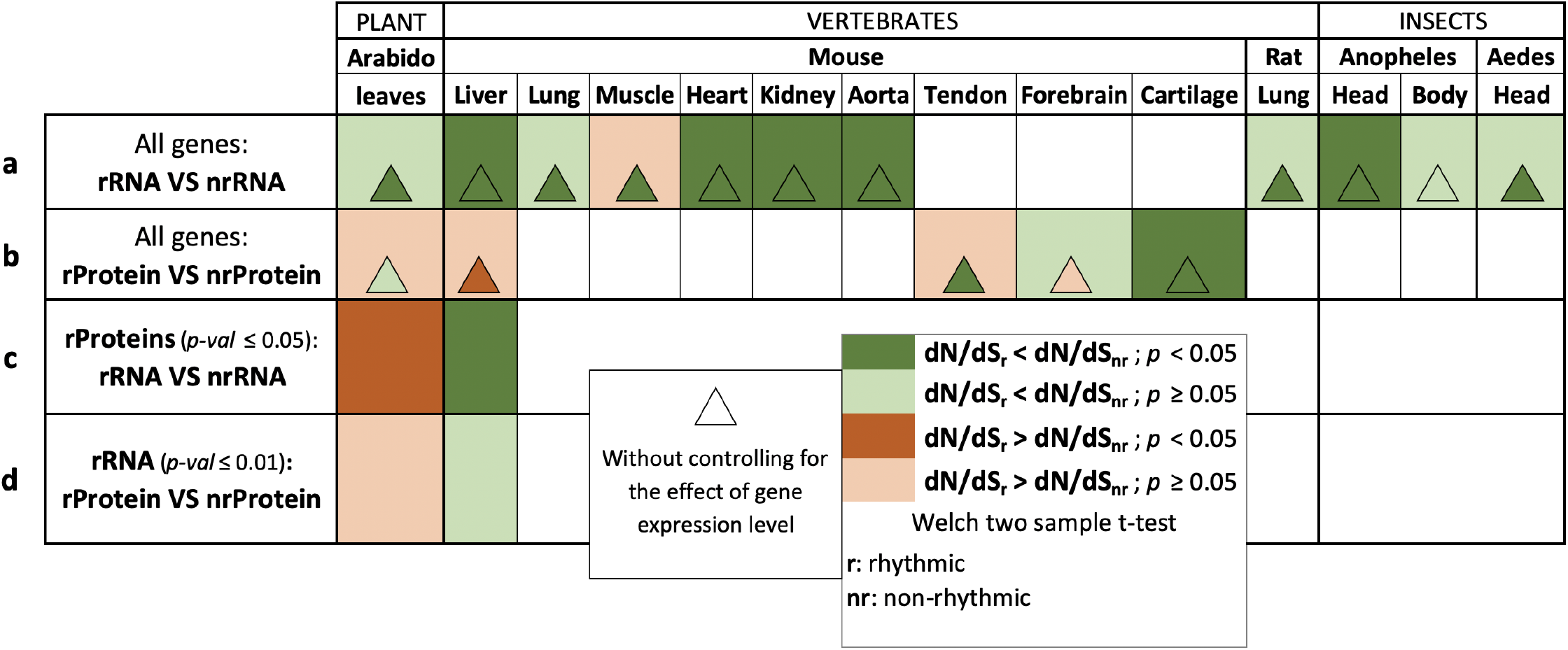
Simplified table showing the results of the Welch two sample t-test testing the hypothesis that dN/dS ratio is equal between rhythmic versus non-rhythmic transcripts (**a**), or proteins (**b**), or between rhythmic versus non-rhythmic transcripts among rhythmic proteins (**c**), and between rhythmic versus non-rhythmic proteins among genes with rhythmic transcripts (**d**). Triangles give the result of the Welch two sample t-test without controlling for the effect of gene expression level. Complete results are provided in Supplementary Table S6.

## Discussion

In this work, we investigate and discuss the evolutionary origins of rhythms in gene expression that respond to factors oscillating on a 24-hour time scale. This scale is longer than those of many biological oscillators at the molecular level, which are ultradian; for instance ~3h for the p53-Mdm2 system, or on the order of minutes to seconds for calcium oscillations. Hence, natural selection was probably instrumental in reaching the longer nycthemeral period of gene expression for many genes.

The endogenous generation of circadian rhythms is an anticipation strategy, which is optimized if the internal clock resonates with the external cycle (Ouyang et al. 1998, Dodd et al. 2005). The autonomous nature of such mechanisms provides a clear advantage to the organism able to anticipate its environmental changes before they take place, allowing it to be “ready” before organisms who would not be endowed with such capacity. In terms of adaptability in a given environment, if we consider nycthemeral rhythms without considering their endogenous or exogenous nature, one can ask the question of the evolutionary origin of maintaining large cyclic biological systems, with gene and tissue specific regulation. Indeed, they are very widely found among living organisms, at every level of biological systems, highlighting their necessity for adapted phenotypes.

An important question which we have addressed is why some genes are rhythmic at the RNA but not the protein level, and vice versa. Apart from mechanistic causes that explain `how`, it was not clear `why` in some cases the rhythmicity is initiated early then lost over the processes (from transcription initiation to protein decay), while in others it is initiated later. In some cases, these lost or gained rhythms might be by-products of the evolution of regulatory processes, but it seems unlikely that all nycthemeral genes be due only to genetic drift. Our previous results showing that rhythmicity of gene expression tends to be a conserved property (Laloum & Robinson-Rechavi 2020) support this perspective. Indeed, we have found a stronger conservation signal of rhythmic expression for evolutionarily closer species (Laloum & Robinson-Rechavi 2020). Moreover, it should be noted that rhythmic expression has costs, e.g., of regulation, and thus it is not always obvious that it should be adaptive. For instance, regulatory dynamics can cause substantial changes in noise levels, e.g. the noise strength immediately following gene induction is almost twice the final steady-state value (Thattai & van Oudenaarden 2001).

### Rhythmic proteins: cost optimization

Our calculation of energetic costs is based on the costs that the cell needs to produce a given steady state protein level. The cell also consumes energy to maintain the steady state protein level. We argue that these costs should correlate with our estimation of expression costs (see section 3 in Supporting information). Indeed, the costs of maintaining a steady state protein level are higher for highly expressed than for lowly expressed genes, even if higher expressed proteins have longer half-lives (see section 4 in Supporting information). Thus, even taking into account a more complex cost calculation would not change our observation that rhythmic genes are costlier.

Our results suggest that rhythmicity of protein expression has been favored by selection for cost control of gene expression, while keeping optimal expression levels. In the case of rhythmic genes, what would an optimal constant level be? We can propose two hypotheses. The first is that it would be the mean expression over the period, since this maintains the same overall amount of protein. The second is that it would be the maximum over the rhythm period, since that is the level needed at least at some point. The second hypothesis explains better the existence of this maximum level during the cycle. Of note, it also strengthens the case for selection on expression cost. Thus, for the case of rhythmic genes, the optimal contant level should at least correspond to the mean expression level (Fig 1d). We provide results obtained using both the maximum and the mean of expression in Fig. 2a.

Another hypothesis is that rhythmicity at protein level could have been selected for noise adjustment. If that were the case, we would expect that for a given mRNA level, the peak level of proteins (*i.e.*, the highest translation rate) would be associated with lower noise to maximize the precision when the protein level is optimal. However, both theoretical and empirical arguments contradict this expectation. For a given mRNA level, theoretical models predict that protein level will be negatively correlated with noise (Hausser et al. 2019, Taniguchi et al. 2010) (section 6.2 in Supporting information). Moreover, measurements of cell-to-cell variability of protein level show that for a given mRNA level, changes in protein level have little to no impact on gene expression precision (Hausser et al. 2019) (section 6.2 in Supporting information). Thus, translational efficiency does not seem to be the main driver of expression noise, and selection on noise cannot explain rhythmicity at protein level.

### Rhythmic mRNAs: noise optimization

Relatively to translational noise, transcriptional noise is the main source of the overall noise (Raj & van Oudenaarden 2008) (section 6.1 in Supporting information). Small fluctuations of expression away from the optimal wild-type expression has been shown to impact organismal fitness in yeast, where noise is nearly as detrimental as sustained (mean) deviation (Schmiedel et al. 2019). Furthermore, noise-increasing mutations in endogenous promoters have been found to be under purifying selection (Metzger et al. 2015), and noise has been shown to be under selection in fly development (Liu et al. 2020). Thus, since we find lower noise among rhythmic transcripts, rhythmic expression of RNAs might be a way to periodically reduce expression noise of highly expressed genes (Figure 2 and Fig. S1), which are under stronger selection. Indeed, we found that genes with rhythmic transcripts are under stronger selection, even controlling for expression level effect. As proposed by Horvath et al. (2019) and supported by results in mouse by Barroso et al. (2018), genes under strong selection could also be less tolerant to high noise of expression. Thus, periodic accumulation of mRNAs might be a way to periodically reduce expression noise of noise-sensitive genes (Fig 1c), *i.e.* genes under stronger selection. However, our results are limited by the fact that noise estimation is based on a single time-point measurement since no scRNA time-series data are currently available for these species. Since the peak time of rhythmic transcripts is distributed across all times (Supporting Information Fig. S9a), the mean noise estimated at a given time-point includes the noise of the genes that are peaking at that time (lowest noise) and all the others that have a higher noise than those at their own peak time-point (Supporting Information Fig. S9b). Our results suggest that rhythmic genes peaking at the time-point of the scRNA measurement have sufficiently low noise for the mean noise of rhythmic genes to be much lower than that of non-rhythmic genes. We suspect that, over a gene-specific threshold of mRNA level, the likelihood that ribosomes interact with mRNA molecules increases much faster than under this threshold, allowing to maximize expression accuracy at the peaks. Future experiments would allow supporting these suggestions.

Studies show that in a population of unicellular organisms, variation in gene expression among genetically identical cells produces heterogeneous phenotypes conferring selective advantages in stressful, changing, or fluctuating environments (as shown in *S. cerevisiae* and *E. coli*) (Acar et al. 2008, Wolf et al. 2015, Thattai & van Oudenaarden 2004, Schmutzer & Wagner 2020, Duveau et al. 2018, Liu et al. 2015). The heterogeneity in genetically identical cells ensures that some cells are always prepared for changes of the environment, called “blind anticipation” by Acar et al. (2008). Thus, expression noise would allow phenotypic diversity and improve fitness in fluctuating environments (Urchueguía et al. 2019). We propose that periodic accumulation of transcripts would allow molecular phenotypes to alternate between a diversified state and an optimal one. Thus, the rhythmicity of mRNA or protein levels would allow to benefit both from the control of expression costs and precision required for protein function, and from the expression variability which improves adaptation in a fluctuating environment. Furthermore, the stochasticity of gene expression could also be beneficial to maintain an oscillator system that would be damped otherwise, according to deterministic simulations (Geva-Zatorsky et al. 2006). *I.e.*, the likelihood that ribosomes interact with mRNA molecules decreases faster than mRNA levels, allowing to amplify the rhythmic signal (without considering the translation factors). Thus, rhythmicity of transcripts might lead to periodic stochasticity that maintains the amplitude of oscillations. Our results in mouse are consistent with all of these considerations (Table 1 and Supplementary Table S5), although it was not fully the case for Arabidopsis (Supplementary Table S5). However, this last point might be explained by the tissue-specificity of rhythmic gene expression. Indeed, for Arabidopsis, the time-series dataset come from leaves whereas single-cell RNA data come from roots. In mouse, tissue muscle gave opposite result, possibly because skeletal muscle is one of the most un-rhythmic tissues in the body. Finally, noise data are samples from a single time-point, and therefore do not give access to the potential rhythmic variation in noise.

We propose here evolutionary explanations of nycthemeral rhythms in gene expression, and show their support by data at different levels and from different species. Rhythmic expression of proteins appears to have been selected for cost optimization, whereas that of mRNAs appears to have been selected for noise adjustment, switching between optimal precision and higher stochasticity. Higher precision allows to maximize the robustness of gene expression when the function is most needed, while higher stochasticity allows to maintain oscillations and to present diverse molecular phenotypes. Alternating between these two states, enabled by rhythmicity at the mRNA level, might have improved the fitness of organisms in fluctuating environments. We also found that rhythmic expression involves highly expressed, tissue-specific genes, consistent with the observation of tissue-specific rhythms. Finally, since genes under strong selection could also become less tolerant to high expression noise levels, our results suggest that rhythmicity at the mRNA level might have been under strong selection for these noise-sensitive genes.

## Materials Methods

### Datasets

Datasets details are available in Supplementary Table S1.

#### Mus musculus

Mouse liver transcriptomic and proteomic time-series datasets come from Mauvoisin et al. (2014). Original protein counts dataset was downloaded from ProteomeXchange (PXD001211) file Combined_WT, and cleaned from data with multiple Uniprot.IDs affectations between raw peptides data. Time-serie transcriptomic data was downloaded from the National Center for Biotechnology Information (NCBI) Gene Expression Omnibus (GEO) accession (GSE33726) (Jouffe et al. 2013) Multi-tissues time-serie transcriptomic data used for tissue-specificity expression comparisons are microarray data from Zhang et al. (2014) downloaded from the NCBI GEO accession GSE54652 (see Materials of Laloum and Robinson-Rechavi (2020) for more details). Tissues analysed are: adrenal gland, aorta, brain stem, brown adipose, cerebellum, heart, kidney, liver, lung, muscle, and white adipose. Multi-tissues time-serie proteomic data come from: Mauvoisin et al. (2014) for the liver, Noya et al. (2019) for the forebrain, Chang et al. (2020) for the tendon, and Dudek et al. (2019) for the cartilage. Finally, single-cell data of thousand of cells in organs for which we had time-series datasets, *i.e.* liver, lung, kidney, muscle, aorta, and heart, were downloaded from figshare using R objects from FACS single-cell datasets (Consortium 2018*a*). We kept cellular sub-types which were found in at least 100 cells and that we considered to be characteristic of the tissue. This leads to keep: `endothelial cell of hepatic sinusoid` and `hepatocyte` for the liver; `epithelial cell of lung, lung endothelial cell`, and `stromal cell` for the lung; `endothelial cell, mesenchymal stem cell`, and `skeletal muscle satellite cell` for the muscle; `cardiac muscle cell, endocardial cell`, and `endothelial cell` for the heart; `endothelial cell` for the aorta; `endothelial cell, epithelial cell of proximal tubule`, and `kidney collecting duct epithelial cell` for the kidney.

#### Arabidopsis thaliana

Leaves time-series proteomic data are the Dataset I from Krahmer et al. (2019), cleaned from data with multiple protein identifications. Leaves time-series transcriptomic data were downloaded from the NCBI GEO accession (GSE3416) (Bläsing et al. 2005). Single-cell data of twenty root cells were downloaded from the NCBI GEO accession (GSE46226) (Efroni et al. 2015).

#### Ostreococcus tauri

Unicellular alga proteomics time-series dataset (normalized abundances) come from Noordally et al. (2018. Transcriptomic time-series dataset come from Monnier et al. (2007), was downloaded from the NCBI GEO accession (GSE16422) and was cleaned for genes with too much missing values (more than seven).

#### Synechococcus elongatus (PCC 7942)

Unicellular cyanobacterium proteomics time-series dataset come from Guerreiro et al. (2014). Transcriptomic time-series dataset come from Ito et al. (2009) and was recovered from Guerreiro et al. (2014).

#### Drosophila melanogaster

Transcriptomic time-series datasets for the body, the head, and the heart, come from Gill et al. (2015) and were downloaded from the NCBI GEO accession (GSE64108) (see Materials of Laloum and Robinson-Rechavi (2020) for more details).

#### Papio anubis [Olive baboon]

Multi-tissues time-serie transcriptomic data used for tissue-specificity expression comparisons are RNA-seq data from Mure et al. (2018) (see Materials of Laloum and Robinson-Rechavi (2020) for more details).

### Pre-processing

For each time-series dataset, only protein coding genes were kept. ProbIDs assigned to several proteins were removed. Probesets were cross-referenced to best-matching gene symbols by using either Ensembl BioMart software (Zerbino et al. 2017), or UniProt (Consortium 2018*b*).

### Rhythm detection

To increase power of rhythm detection (Laloum & Robinson-Rechavi 2020), we considered biological replicates as new cycles when it was possible. We used `GeneCycle` R package (version 1.1.4) (Ahdesmaki et al. 2012) available from CRAN and used the robust.spectrum function developped by (Ahdesmäki et al. 2005) with parameters *periodicity.time=24* and *algorithm=”regression”* - that computes a robust rank-based estimate of the periodogram/correlogram and that we improved with the `try-catch` function to avoid error of dimension with MM-estimation method. When the *p*-values distribution obtained did not correspond to the expected distribution - skewed towards low *p*-values because of the presence of rhythmic genes - we used the results obtained by the rhythm detection method used by the original paper from where the data came after checking they presented a classic skewed *p*-values distribution. Finally, for each gene or protein having several data (ProbIDs or transcripts), we combined *p*-values by Brown’s method using the `EmpiricalBrownsMethod` R package (See Supporting information). Thus, for each dataset, we obtained a unique rhythm *p*-value per gene or per protein. Due either to low-amplitude or less-accurate measurements, it was more challenging to identify rhythms in proteomics data. That is why, in general, we used lower stringency for proteins.

### Consistent gene expression levels

Tissue-specific mRNA or protein abundances were the average or the maximum level of the *n* time-points such as for gene *i*:

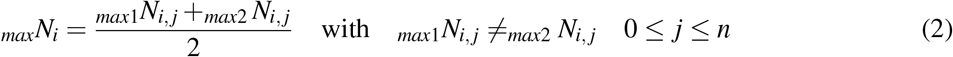

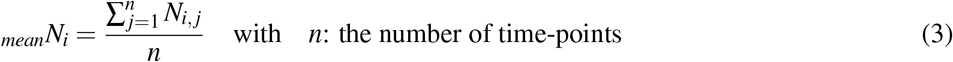

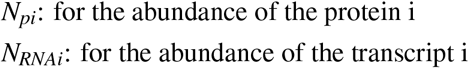

### Expression costs

Energetic costs of each AA (unit: high-energy phosphate bonds per molecule) come from Akashi and Gojobori (2002), or Wagner (2005) which are linearly correlated (Fig. S5). The averaged AA synthesis cost of one protein of lenght *Lp* is:

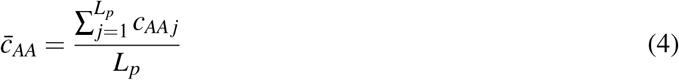

Protein sequences come from FASTA files downloaded from EnsemblPlants (Zerbino et al. 2017) or UniProt (Consortium 2018*b*) for: Proteome UP000002717 for *Synechococcus elongatus* PCC 7942, Uniprot Reviewed [Swiss-Prot] for *Mus musculus*, Proteome UP000009170 for *Ostreococcus tauri*.

Main results use Wagner (2005) AA costs data. We provide supplementary results using Akashi and Gojobori (2002) AA costs data (Fig. S6).

### Multi-tissues analysis

To obtain comparable expressions levels between different tissues or datasets, we normalized expression values by Z-score transformation such as in the dataset of *n* genes, the mean expression of the gene *i* becomes:

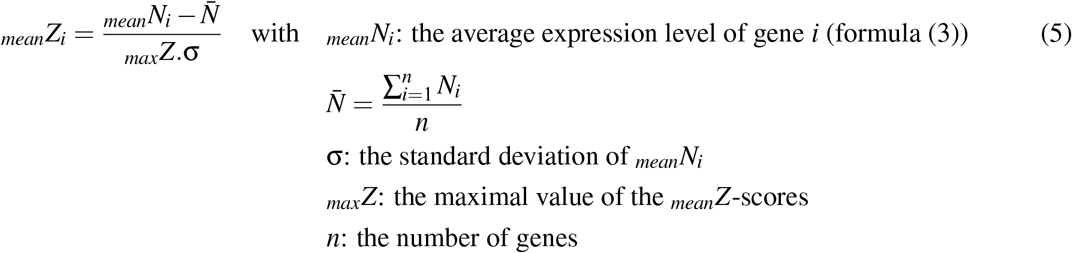

and the maximal expression of the gene *i* becomes:

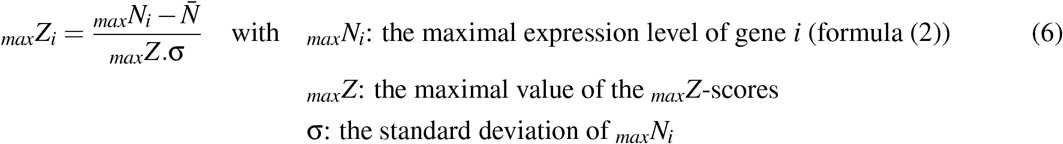

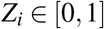

To compare the expression levels between the set of *n*_*r*_ rhythmic tissues and the set of 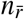 non-rhythmic tissues, we estimated the difference (δ) of expression levels between these two groups for each gene *i* such as:

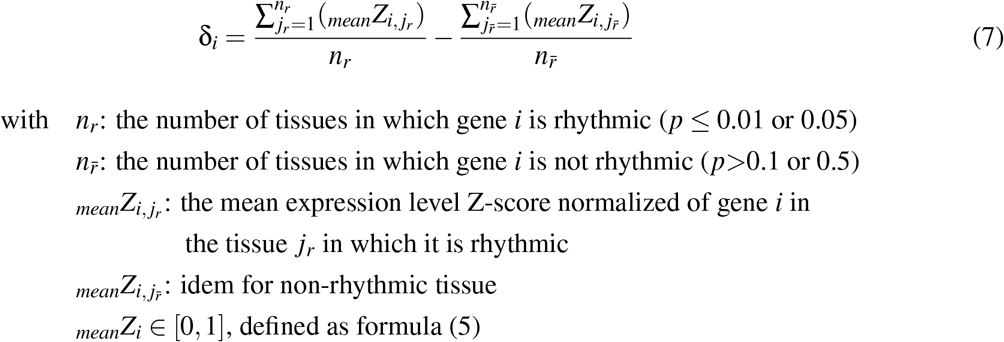

Finally, we analysed the distribution of δ_*i*_ and generated a Student’s t-test to compare its mean distribution with an expected theoretical mean of 0.

### Tissue-specificity of gene expression

To calculate a tissue-specificity τ for each gene *i*, we log-transformed the averaged gene expression and followed Kryuchkova-Mostacci and Robinson-Rechavi (2016) to make expression values manageable. Thus, among the *n* tissues, the tissue-specificity of gene *i* is:

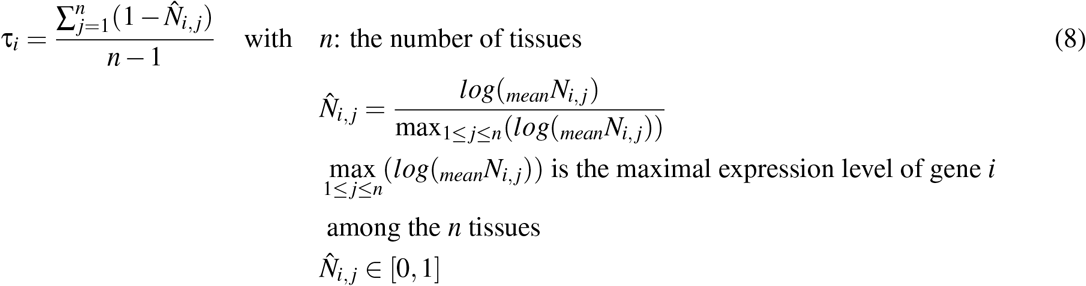

This tissue-specificity formula was described by Yanai et al. (2004). Finally, we performed linear regressions to analyse the respective and the interaction influences of gene expression level and tissue-specificity into the rhythmicity. For mouse (Zhang et al. 2014), we used RNA-seq data to estimate the mean expression used for the calculation of τ, and used rhythm *p*-values obtained from microarray dataset since there was more time-points in the microarray time-series (as discussed in benchmark paper). Figure S7 shows the distributions of τ obtained.

### Gene expression noise quantification

We estimated a unique expression noise per gene by taking advantages of *Arabidopsis thaliana* roots (Efroni et al. 2015) and Mus musculus liver, lung, kidney, muscle, aorta, and heart (Consortium 2018*a*) (from figshare using R objects of FACS single-cell datasets) single-cell RNAseq data (see Materials). For Arabidopsis, we obtained gene expression for twenty root quiescent centre (QC) cells for which a total of 14,084 genes had non-zero expression in at least one of the 20 single cells. For mouse, we obtained gene expression for a number of cells ranging from 180 to 3772 cells, for which a total of genes with non-zero expression was around 19,000 genes. As in Barroso et al. (2018), we kept only genes whose expression level satisfied log[FPKM + 1] *>*1.5 in at least one single cell. We calculated the stochastic gene expression (*F*\*) defined by Barroso et al. (2018) as a measure for gene expression noise (cell-to-cell variability) designed to control the biases associated with the correlation between the expression mean (*μ*) and the variance (σ^2^) by using the lowest degree polynomial regression which decorrelate them. We tested different degrees of the polynomial regression to estimate log(σ^2^) in the calculation of *F*\* and measured the correlation based on Kendall’s rank and linear regreassion slope (see Supporting information).

### dN/dS analysis

dN/dS data have been downloaded from Ensembl BioMart Archive 99 (Zerbino et al. 2017). The homologous species used are: *Mus musculus* - *Rattus norvegicus*; *Arabidopsis thaliana* - *Arabidopsis lyrata*; and *Anopheles gambiae* - *Aedes aegyti*. We first tested the hypothesis that rhythmic (*p*-value≤cutoff.1) versus non-rhythmic genes (*p*-value*>*cutoff.2) have equal ratio of non-synonymous to synonymous substitutions. Then, since rhythmic genes are now known to be enriched in highly expressed genes and because highly expressed genes are under purifying selection, we controlled for the effect of gene expression on dN/dS ratio by testing the hypothesis that rhythmic (*p*-value≤cutoff.1) versus non-rhythmic (*p*-value*>*cutoff.2) genes among residuals of the linear regression fitting gene expression level and dN/dS ratio. Complete results are given in Supplementary Table S6 and in a simplified way in Table 2.

In R code it gives:

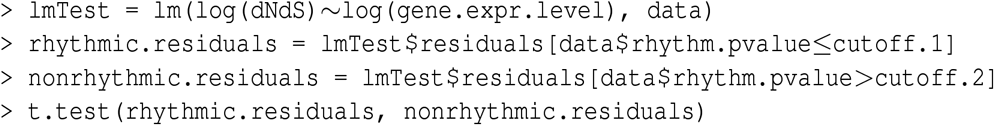

Plots have been generated using ggplot2 package (version 3.3.2) run in R version 4.0.2.

## Availability of data and scripts

Data and scripts are available at: https://github.com/laloumdav/cost_noise_conservation_rhythmicity

## Acknowledgements

We thank Johanna Krahmer (CIG, UNIL, Lausanne Switzerland) for her helpful advice on datasets in plants. This work was supported by Swiss National Science Foundation grant 31003A 173048 and by University of Lausanne.

## Conflict of Interests

The authors declare that they have no conflict of interest.

## Notes

https://github.com/laloumdav/cost_noise_conservation_rhythmicity

### Competing Interest Statement

The authors have declared no competing interest.

### Summary of Updates

Revision following reviews at Reviews Commons.

