## Supplementary text for "Two levels of selection of rhythmicity in gene expression: energy saving for rhythmic proteins and noise optimization for rhythmic transcripts"

### Supporting information
Figures S8-S20

### Energetic costs and expression noise of rhythmically expressed genes

#### 1 Detection of rhythmic gene expression

We mainly used the algorithm GeneCycle to detect rhythmic patterns in gene expression time-series (See Methods for more details). We checked that density distributions of $p$-values obtained from rhythm detection methods used in this paper produced expected left-skewed distributions (Fig. S10 and S12). For each gene or protein with several data (several ProbIDs or transcripts), we combined $p$-values by Brown's method using the `EmpiricalBrownsMethod` R package. Figure S11 shows the density distribution of $p$-values obtained after this Brown's normalization for the transcriptomic data in Ostreococcus.

#### 2 Gene expression level

Figures S13 and S14 show that distributions of the mean (Fig S13) and the maximum (Fig S14) expression level calculated over time-points (See Methods) can be considered as normally distributed, so relevant for our statistical analysis that need these initial conditions.

#### 3 Gene expression costs

To produce a given steady state protein level $N_p$, the cell request energetic costs at transcriptional level ($C_{RNA}$) and at translational level ($C_p$):

$$C_{RNA} = N_{RNA}.L_{RNA}.\bar{c}_{nt} \quad (1)$$

$$C_p = N_p.L_p.\bar{c}_{AA} \quad (2)$$

$N_{RNA}$: abundance of transcripts
$L_{RNA}$: transcript length
$\bar{c}_{nt}$: averaged nucleotide synthesis cost of the transcript
$N_p$: abundance of proteins
$L_p$: protein length
$\bar{c}_{AA}$: averaged AA synthesis cost of the protein

The number of ATP molecules consumed is in averaged around 30∼P for one amino-acid (AA) molecule produced (estimated in *E. coli*) (Wagner 2005), against 49∼P for one single nucleotide (estimated in yeast and *E. coli*) (Wagner 2005, Lynch & Marinov 2015). The median length is estimated around 1500 nucleotides for transcripts against 400 AA for proteins in yeast (Wagner 2005, Lynch & Marinov 2015). And, the abundance of transcripts is in average 1.2 molecules per cell (Wagner 2005), against 2,622 proteins per cell (yeast) (Ho et al. 2018) (9 and 6,384 molecules respectively found recently by Lahtvee et al. 2017). Thus, even whether the precursor synthesis per nucleotide is 1.6 fold costlier than for AA molecule and proteins are around 3.75 fold shorter than transcripts, $N_p/N_{RNA}$ is on the order of 1,000, so the cost at the protein level is 160 times that at the RNA level (even higher at slower cellular growth-rate). Finally, protein costs are even more higher since chain elongation costs are larger than those of nucleotides polymerization since the nucleotide precursors are already activated molecules (rNTP) whereas chain elongation needs costs of charging tRNAs with AA (Lynch & Marinov 2015).

Thus, because $C_{RNA} << C_p$, $C_p$ gives an estimate broadly representative of the expression costs per gene. Finally, since we compared these costs between genes, the other costs such as degradation or chain elongation costs, should not change our results. Degradation may cost essentially nothing (Wagner 2005) and its cost must be correlated with $C_p$ since longer or higher expressed proteins need more chain elongation activity and degradation rate (see next section), so does not modify the comparison of expression costs between genes.

Our results support the hypothesis also claimed by Wang et al. 2015 that cycling expression of the more expensive genes is a conserved strategy for minimizing overall cellular energy usage. In this study, we provide new results based on relevant data. Indeed, data used by Wang et al. 2015 for the calculation of costs appears to be biased, partly because: i. translation rates come from fibroblasts cells (Schwanhäusser et al. 2013); and ii. there were errors in the estimation of protein levels resulting in a systematic underestimation of protein levels and derived translation rate constants (Cf Corrigendum (Schwanhäusser et al. 2013)).

### 4 Expression costs, decay and half-lives

#### 4.1 Costs of protein decay

Protein half-life is the time requested to reduce the protein amount by 50%. In terms of protein concentration, assuming constant protein production, protein half-life depends on protein degradation and cell dilution (cell's growth) (equation (3) and (4)). We assume that spontaneous degradation of proteins is negligible.

$$T_{1/2} = \frac{ln(2)}{\delta} \tag{3}$$

$T_{1/2}$: protein half-life
$\delta$: protein decay

$$\delta = \delta_{sdeg} + \delta_{deg} + \delta_{dil} \tag{4}$$

$\delta_{sdeg}$: spontaneous decay of unstable molecules
$\delta_{deg}$: active protein degradation rate
$\delta_{dil}$: cellular dilution

Costs of protein decay are negligible enough to not be opposed by selection. Indeed, Lynch and Marinov (2015) and Wagner (2005) have shown that "degradation in a lysosome may cost essentially nothing, and amino-acid export back to the cytoplasm consumes ~1 ATP for every 3 to 4 amino acids". Compared with the unique cost of producing one single nucleotide which consume 49~P, protein decay costs becomes negligible comparatively to transcriptional costs, which are themselves negligible comparatively to translational costs. All the more, given that amino acids from degradation are reused and do not need to be produced by the cell, which therefore economizes around 30 ~P per amino-acid.

#### 4.2 Half-lives of rhythmic proteins

In rapidly growing bacterial cells, dilution is often more significant than degradation (Eden et al. 2011, Ingalls 2013), whereas in no-dividing and slowly dividing cells such as differentiated mammalian cells, protein half-life rely mostly on degradation because dilution is negligible (Eden et al. 2011). Half-life might be an inappropriate parameter regarding rhythmic proteins. Indeed, half-life depends on protein production rate and degradation rate which, for rhythmic proteins, vary over time. We think that protein half-life should be a constant property of proteins and, therefore, is difficult to understand for rhythmically expressed genes (although it is discussed by Lück et al. 2014).

Suppose we can deal with protein half-life property for all genes. Since higher expressed genes have longer half-lives (they are more stable) (Martin-Perez & Villén 2017, Mauvoisin et al. 2014) and because highly expressed genes tend to be rhythmic, the expression costs for maintaining high level of these proteins should be reduced (due to their longer life-span). That is why, the cost per time-unit of maintaining the steady-state protein level might not be correlated with our estimation of expression cost ($C_p$). *I.e.*, rhythmic proteins for which our results suggest that they are rhythmic because they are costlier (and costlier because they are highly expressed) would, in fact, require less energetic costs to be expressed than expected because these protein molecules tend to have longer half-lives (they live longer). On the other hand, models suggest that a longer half-life damps the amplitude of the periodic expression (Lück et al. 2014); as supported by observations showing that non-rhythmic proteins have much longer half-lives than rhythmic ones (in mouse fibroblast) (Mauvoisin et al. 2014). Thus, highly expressed genes should be less rhythmic (due to amplitudes damped by longer half-lives) and therefore, less energy saved over 24 hours. Consequently, the rhythmic expression of costly proteins (as our results suggest) should not involve proteins with longer half-lives. Thus, rhythmic and highly expressed proteins remain costlier for the cell per time-unit than rhythmic and lower expressed proteins. Therefore, our results remain consistent even when considering protein half-lives.

### 5 Averaged AA synthesis cost and protein length

Before applying the t-tests, we checked that averaged AA synthesis costs and the length of proteins were normally distributed in both groups (rhythmic, first 15%, and non-rhythmic) (Fig. S15 and S17). We also verified that quantile-quantile-plots showed comparable distributions between the theoretical distribution and the empirical distribution for both groups (Fig. S16 and S18).

### 6 Gene expression noise

#### 6.1 Transcriptional noise is the main source of the overall noise

Relatively to translational noise, transcriptional noise is the main driver of the overall noise (Raj & van Oudenaarden 2008) and should give a good estimation of the output noise. Indeed, based on estimations of coefficient of variations (CV, cell-to-cell variations of protein level) for diverse transcription and translation rates in *E. coli* and *S. cerevisiae*, Hausser et al. 2019 have shown that for a fixed transcriptional rate, CV is almost constant for diverse translational rates. Thus, changes in protein level have little to no impact on gene expression noise. The availability of mRNA molecules seems to drive the final noise. *I.e.*, comparatively to the noise caused by the translational activity, the availability of low number molecules such as transcriptional factors (subject to the stochasticity of diffusion and binding in the cell environment) is the main factor of the output cell-to-cell variation in protein abundances.

#### 6.2 Rhythmic proteins and expression noise

In the case of genes with constant mRNAs abundances and with rhythmic proteins, we expect that rhythmic expression at the protein level cannot originate from selection on expression noise. Indeed, since increasing transcription at constant protein abundance decreases expression noise (Hausser et al. 2019, Taniguchi et al. 2010) (Fig. S8A), we should theoretically expect to improve expression precision (decrease noise) when protein level decreases for an equi-mRNA level, because the translational efficiency decreases (i.e. less proteins are produced per mRNA molecule). Thus, in this case, we expect the gene expression precision to be highest when the protein level is at the valley and lowest when the protein level is at its peak which we presume to be near to the optimal protein level (Fig. S8B: Theoretical plot). Based on estimations of coefficient of variations (CV, cell-to-cell variations of protein level) for diverse transcription and translation rates in *E. coli* and *S. cerevisiae*, Hausser et al. (Hausser et al. 2019) have shown that for a fixed transcriptional rate, CV is almost constant for diverse translational rates (Hausser et al. 2019). Thus, the translational efficiency might not be the main driver of expression noise (Fig. S8B: last plot). The availability of mRNA molecules drives the final noise, i.e. mainly the availability of transcriptional factors. Thus, this suggest that, in both cases, rhythmicity at protein level cannot be due to a selection on noise.

Finally, we remain attentive to the fact that all these considerations presume an immediacy between expression noise and biological functionality.

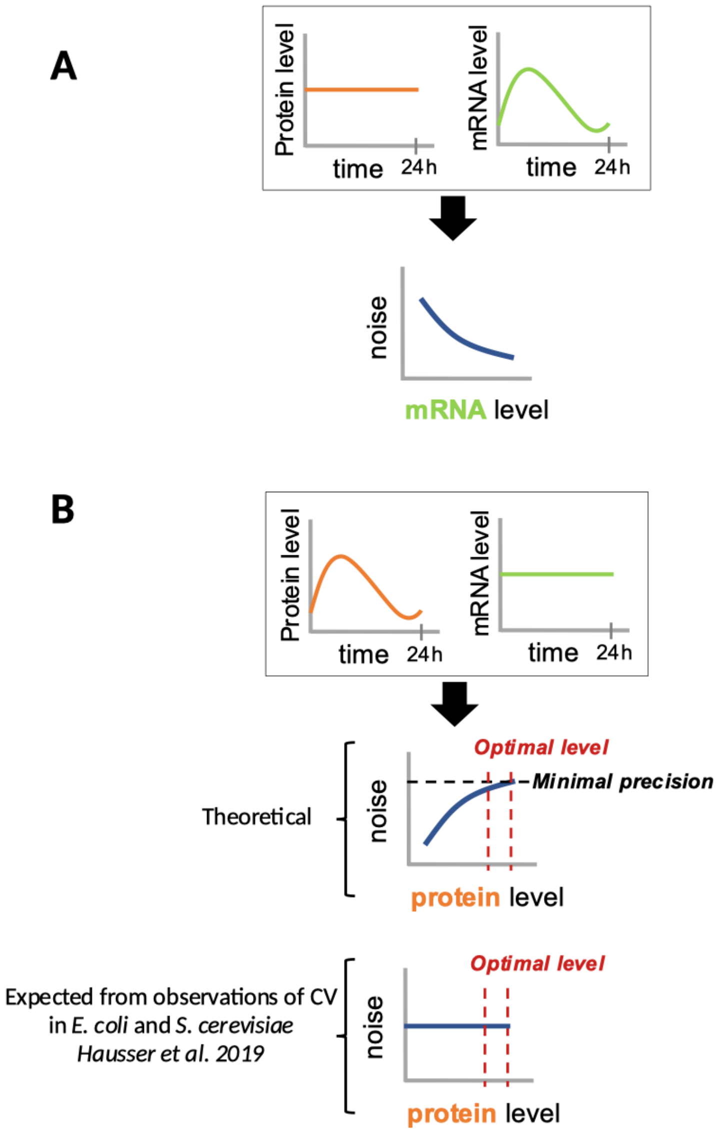


Figure **S8**: **A)** The expression noise decreases with increasing transcription for an equi-protein level (Hausser et al. 2019). In the case of rhythmic mRNAs abundance and constant protein level, the expression noise decreases when mRNAs level reaches its peak. **B)** Theoretical expression noise variation in the case of constant mRNAs abundance and rhythmic proteins. Theoretically, the expression noise is correlated with the translational efficiency. However, observations in *E. coli* and S. cerevisiae show that for a fixed transcriptional rate (that we assume to be correlated with mRNAs level), the expression noise (CV) is almost constant for diverse translational rates (Hausser et al. 2019).

#### 6.3 Noise level is based on a single time-point scRNA dataset

Indeed, our noise estimation is reported at a single time-point (unknown) for a cell population (scRNA). Since the peak times of rhythmic genes is largely distributed (Fig. S9a), we expect the mean noise of a given time-point to be general to all time-points (Fig. S9b).

#### 6.4 Noise estimation

We compared several noise estimation methods and found that the $F*$ polynomial degree of Barroso et al. (Barroso et al. 2018) method was the best method across all datasets and especially the most efficient method in controlling for the effect of mean expression (Supplementary Table S7). To do this, we calculated the slope of the linear model which best fit the correlation between the noise and mean expression (Fig. S19 and S20), as well as the $R^2$ and the Kendall correlation ($\tau$) , applied to normally distributed noise estimations. The linear model that explained the variance the least well (small $R^2$) was considered to be approximately the best expected value. Indeed, we expect a good noise estimation model to be independent of the mean gene expression (linear regression slope and Kendall's $\tau$ near to 0) and with a large range of noise values for every mean expression (small $R^2$). The $F*$ polynomial degree of Barroso et al. (Barroso et al. 2018) was the best method because it removed the correlation between mean expression and $F*$ (smallest linear regression slope and Kendall's $\tau$) and keep a large range of noise values for every mean expression (smallest $R^2$) (Supplementary Table S7). If none of Kendall's correlation was not significantly uncorrelated (all Kendall's $p$-value<0.05), we used the polynomial degree which seem to maximally removed residual correlations (smallest Kendall's $\tau$ and smallest linear regression slope). Thus, we used $F*$: degree 3 for mouse liver (Kendall's $\tau$ = -0.0288, $p$-value = 1.6e-07, linear regression slope = 1.14e-13); degree 3 for mouse lung (Kendall's $\tau$ = -0.0545, $p$-value < 2.2e-16, linear regression slope = 3.40e-15); degree 3 for mouse muscle (Kendall's $\tau$ = -0.0795, $p$-value < 2.2e-16, linear regression slope = 9.76e-15); degree 4 for mouse heart (Kendall's $\tau$ = -0.0445, $p$-value < 2.2e-16, linear regression slope = -3.75e-13); degree 4 for mouse aorta (Kendall's $\tau$ = -0.0183, $p$-value = 4.70e-04, linear regression slope = 4.83e-14); degree 4 for mouse kidney (Kendall's $\tau$ = -0.0528, $p$-value < 2.2e-16, linear regression slope = 1.04e-13); and degree 4 for Arabidopsis roots (Kendall's $\tau$ = 0.0091, $p$-value = 0.1162, linear regression slope = -9.78e-15).

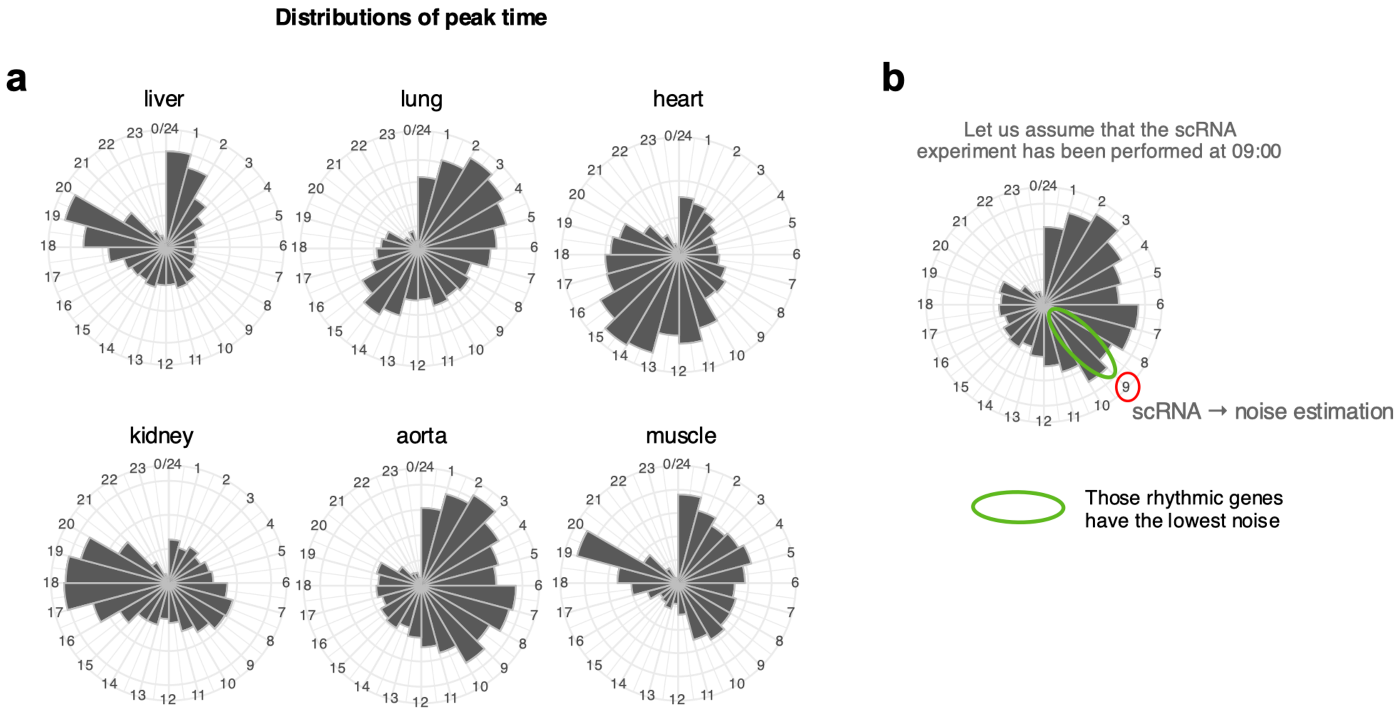


Figure **S9**: **a)** Distributions of peak time for rhythmic transcripts. **b)** Among rhythmic genes, the mean noise estimated at a given time-point includes the noise of the genes that are peaking at that time (lowest noise) and all the others that have a higher noise than those at their own peak time-point. *Hours are not relevant, only the distribution across the 24-hours is.*

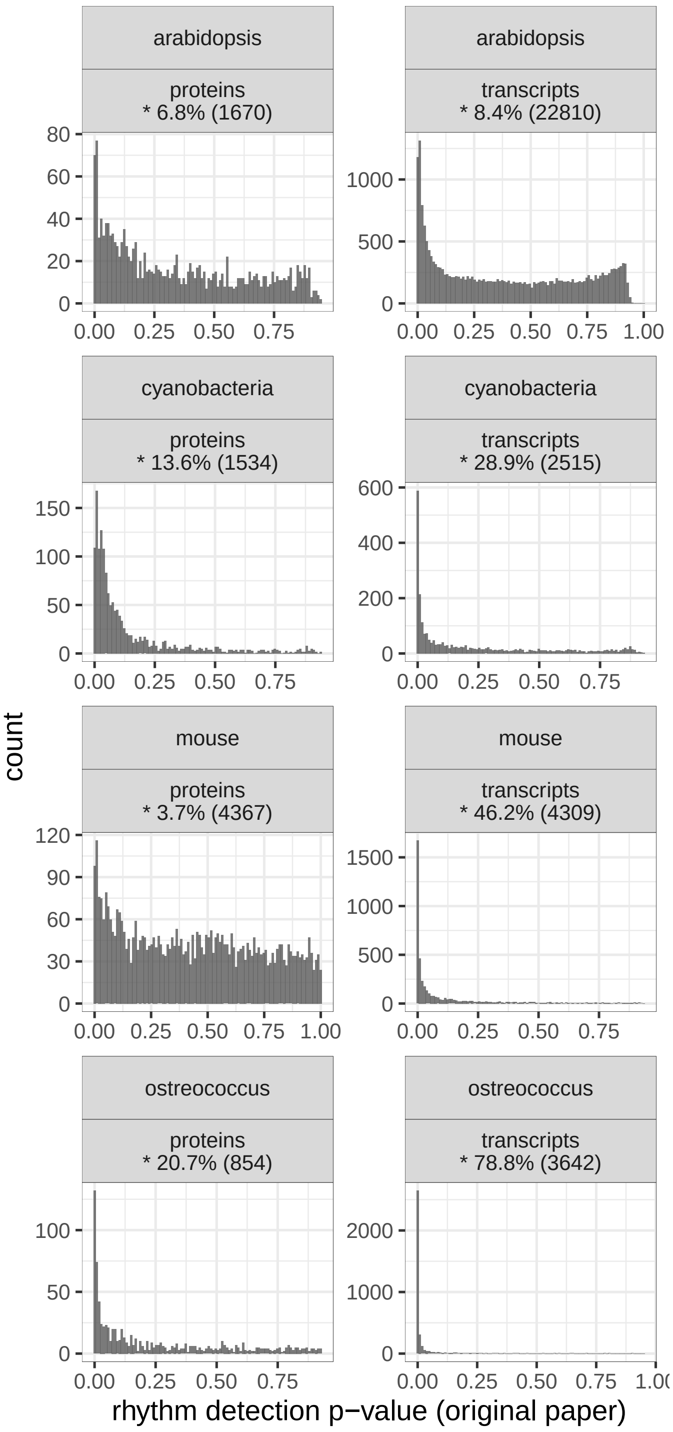


Figure **S10**: **Density distribution of *p*-values obtained from rhythm detection**. All by GeneCycle except for the mouse liver proteome dataset for which no classic rhythm detection methods worked (we used the Harmonic regression method used in the original article). Percentage of genes detected rhythmic with *p*-value$\leq$0.01 (among total of genes).

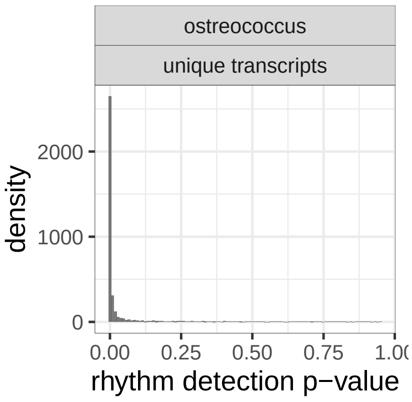


Figure **S11**: Distribution of *p*-values obtained after the Brown normalization to score unique genes in *Ostreococcus tauri* transcripts time-serie dataset.

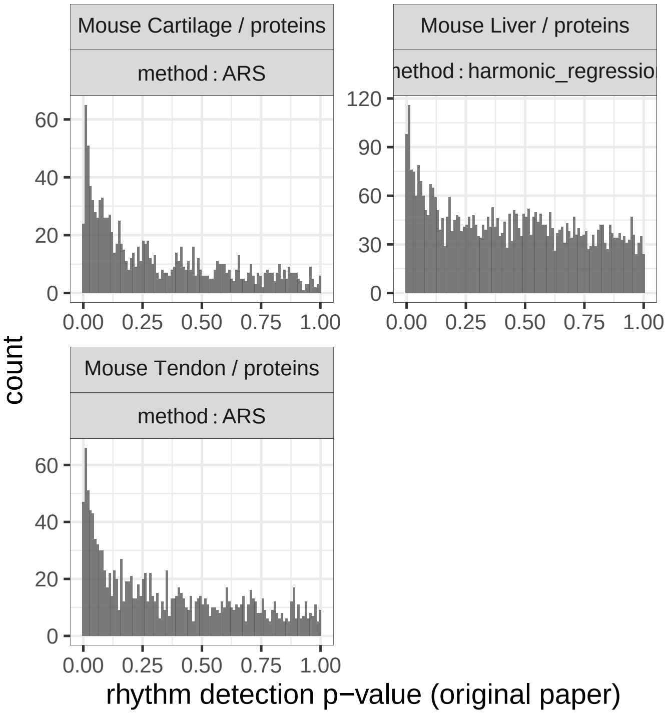


Figure **S12**: Density distribution of *p*-values obtained from rhythm detection using the method used in this paper.

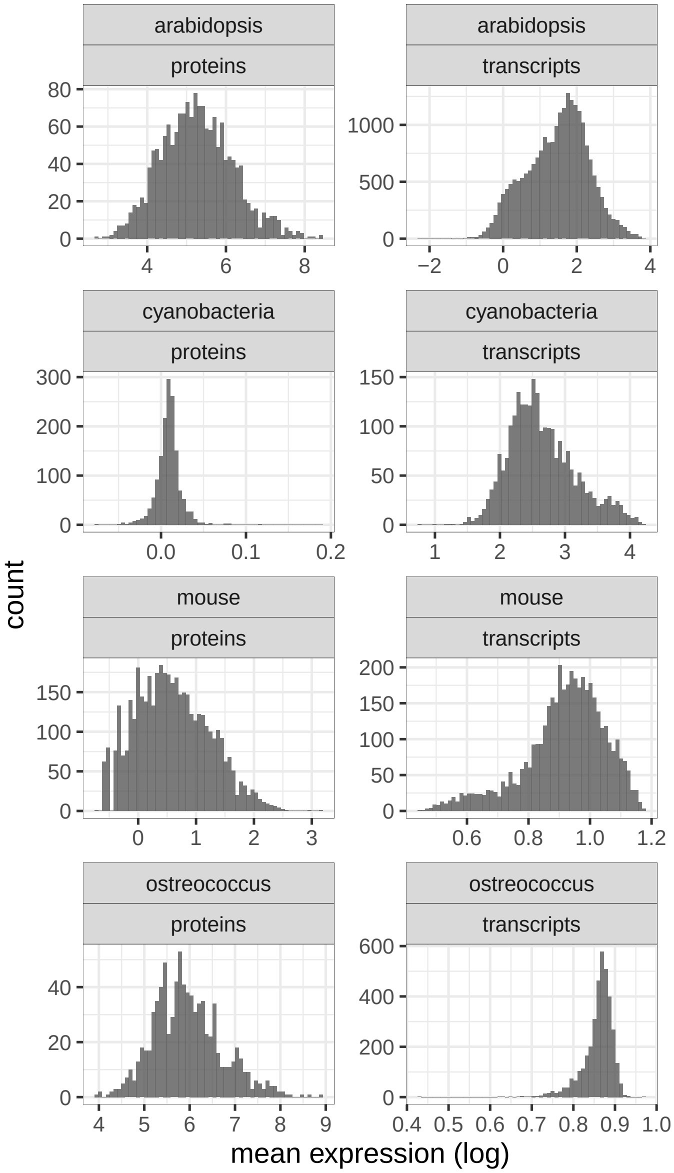


Figure **S13**: Histograms of the mean expression values calculated over time-points.

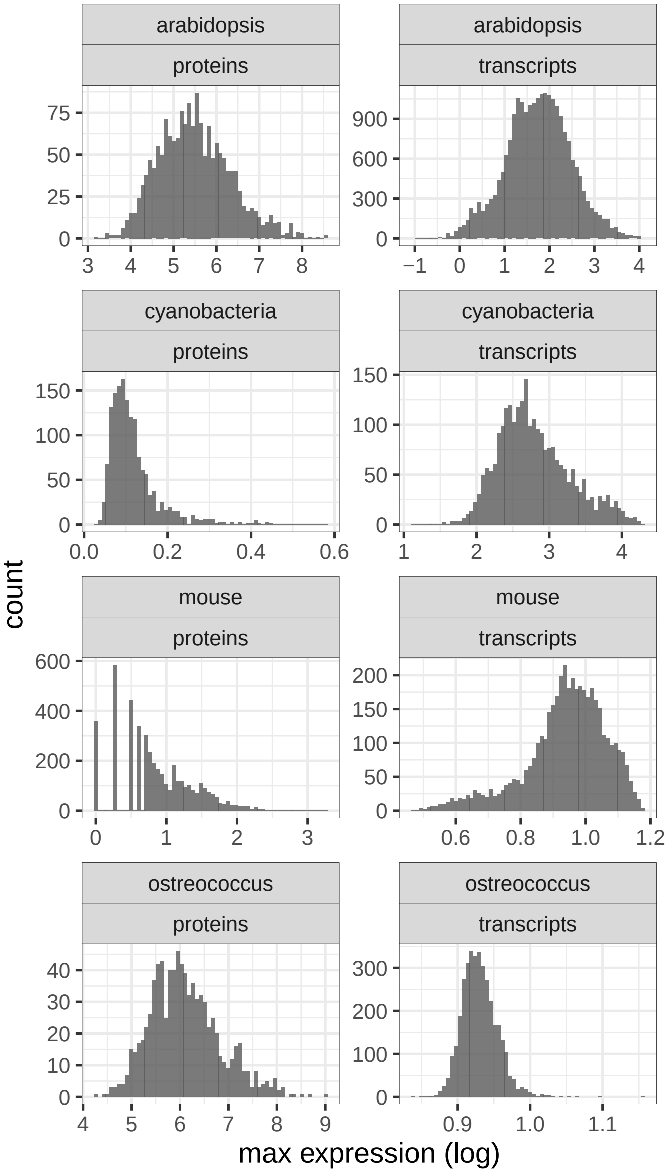


Figure **S14**: Histograms of the maximum expression values calculated (average of the two maximum expression values among time-points).

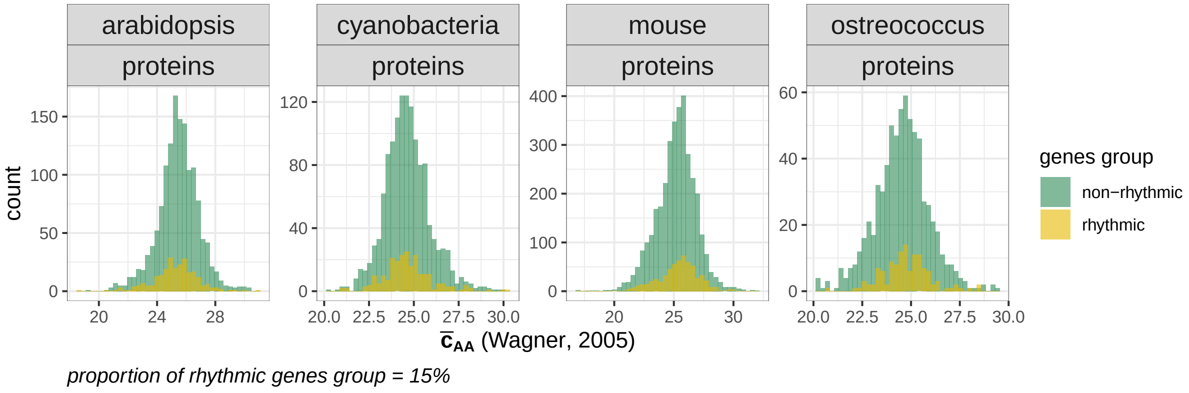


Figure **S15**: Histograms of the averaged AA synthesis costs calculated in both groups.

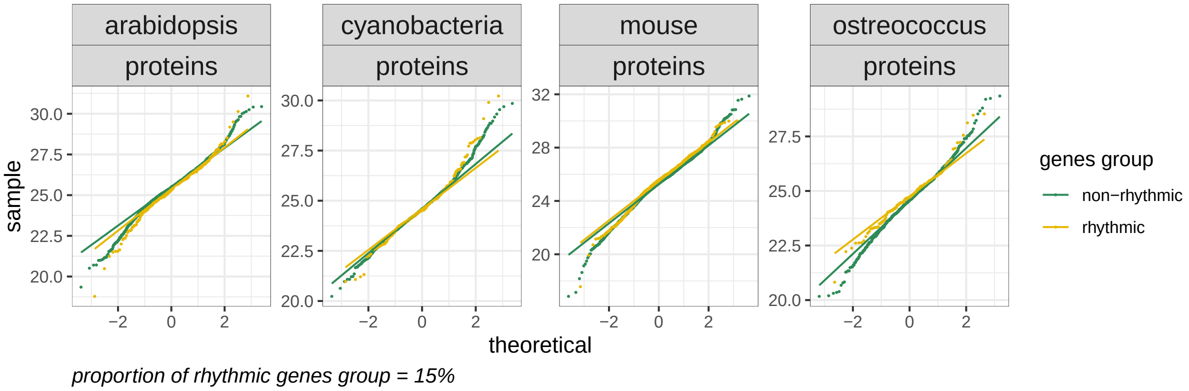


Figure **S16**: QQ-plots of the averaged AA synthesis costs calculated in both groups.

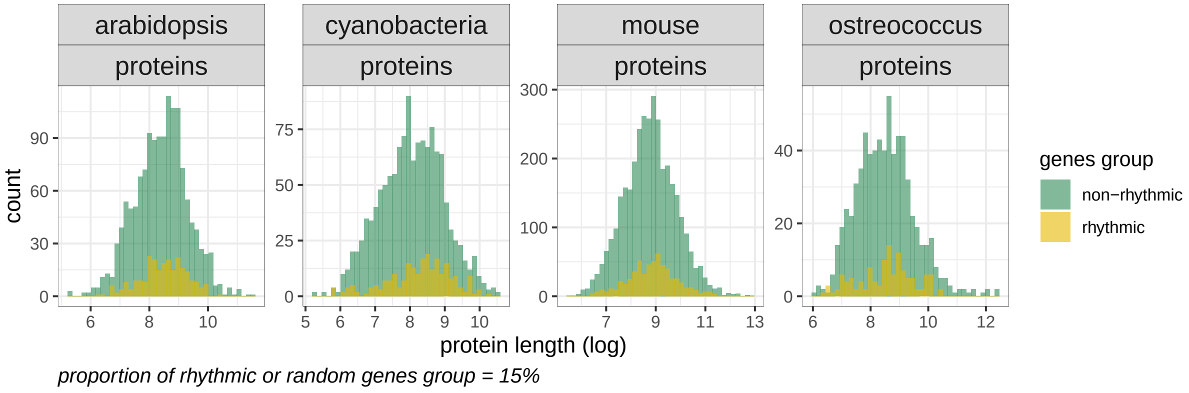


Figure **S17**: Histograms of the protein lengths in both groups.

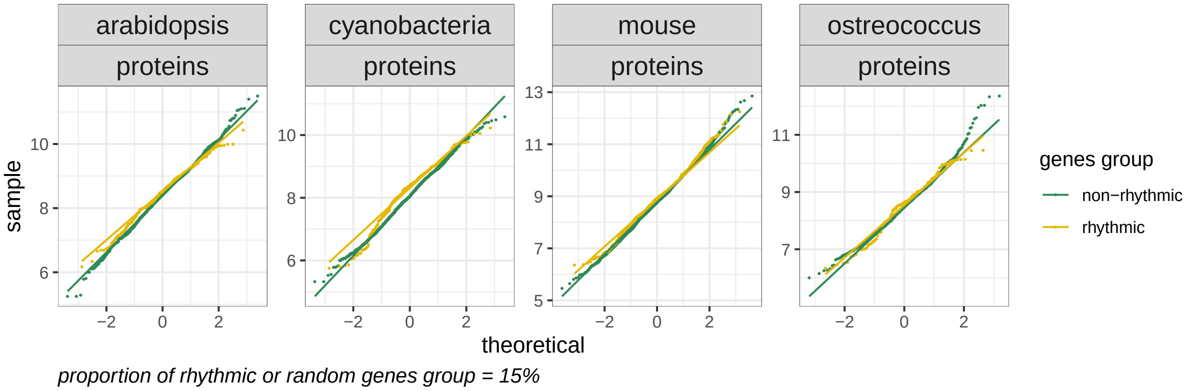


Figure **S18**: QQ-plots of the protein lengths in both groups.

**(a)** ***Arabidopsis thaliana*** **(root)**

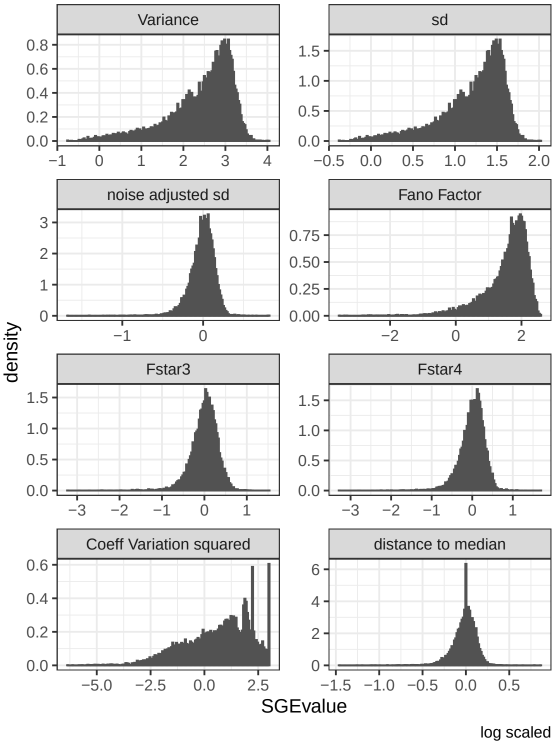


**(b)** ***Mus musculus*** **(Liver)**

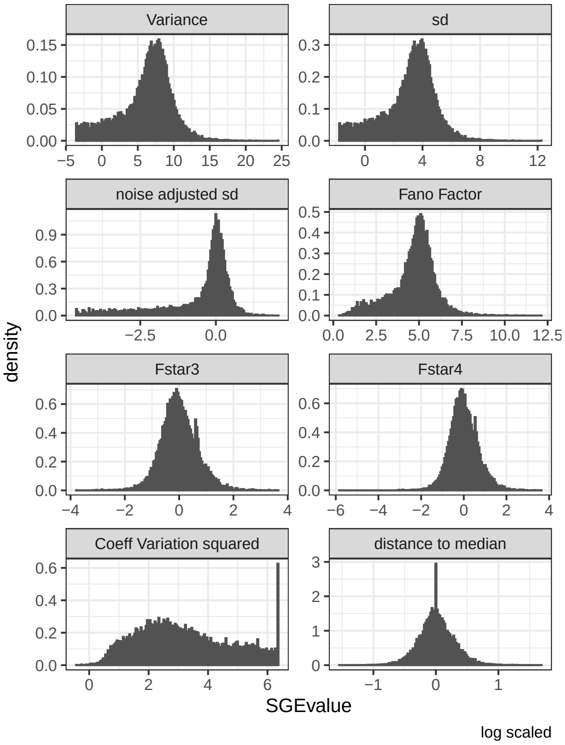


**(c)** ***Mus musculus*** **(Lung)**

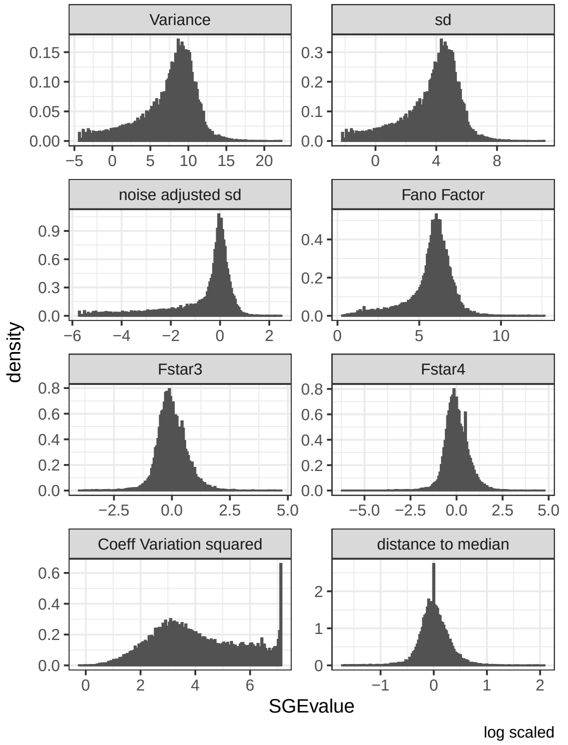


**(d)** ***Mus musculus*** **(Aorta)**

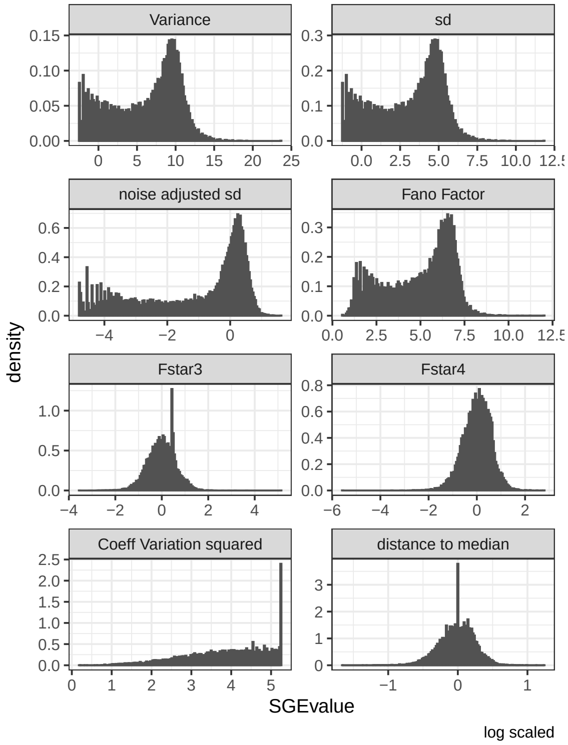

**(e) *Mus musculus* (Heart)**

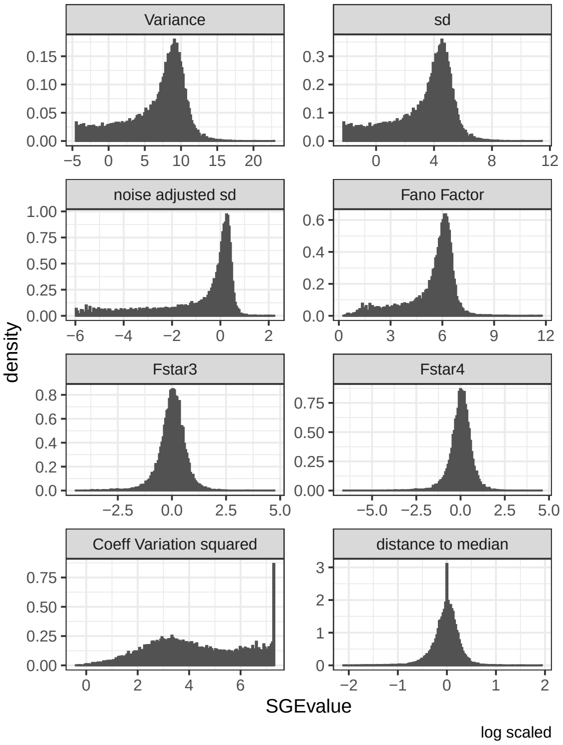


**(f) *Mus musculus* (Kidney)**

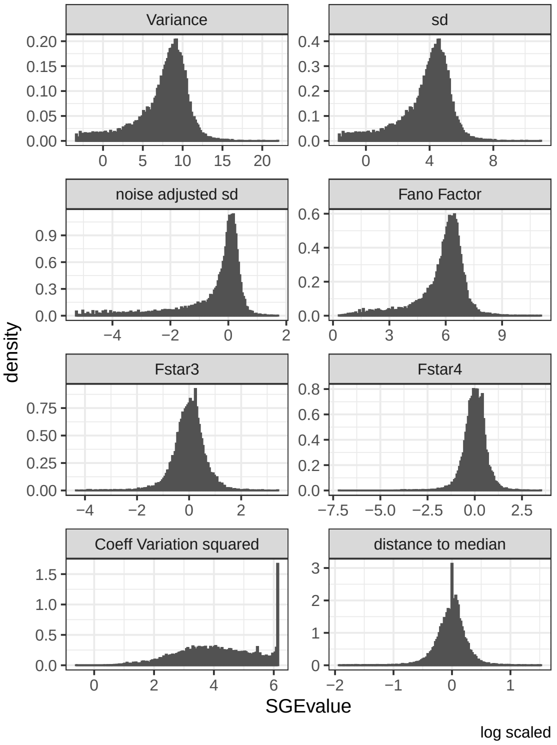


**(g) *Mus musculus* (Muscle)**

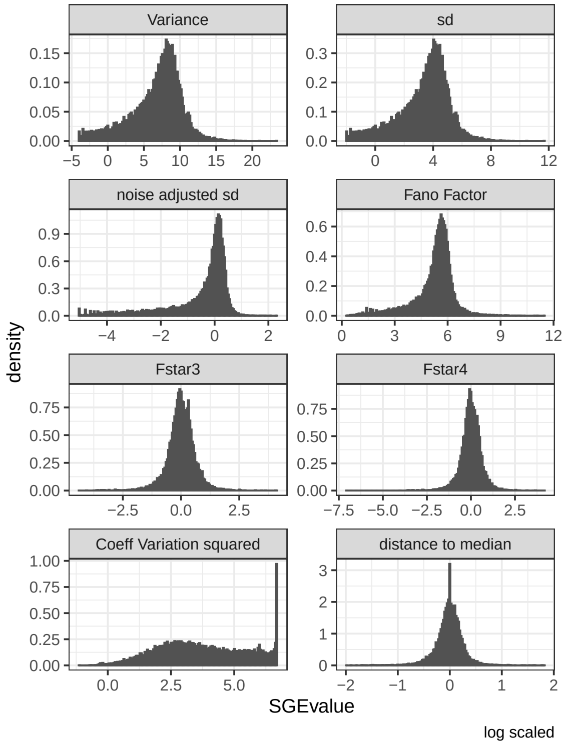


Figure **S19**: **a-f)** Histograms of the variance, standard deviation, and various stochastic gene expression (SGE) estimations. All variables are log transformed.

**(a) *Arabidopsis thaliana* (root)**

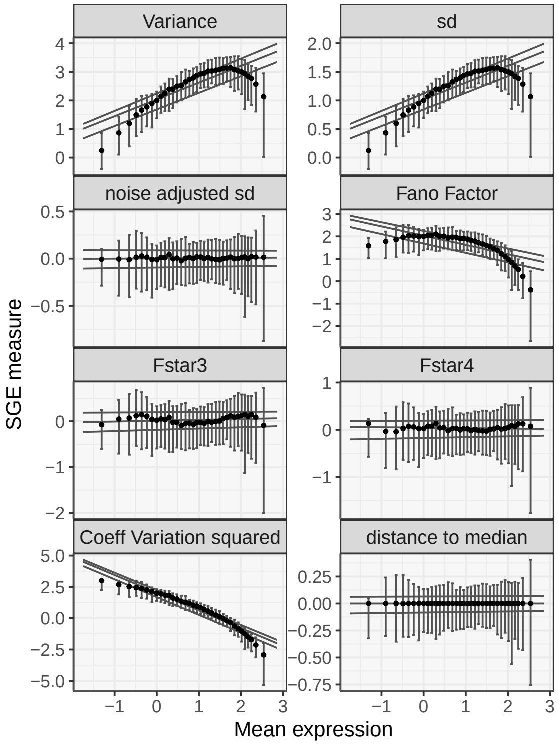


**(c) *Mus musculus* (Lung)**

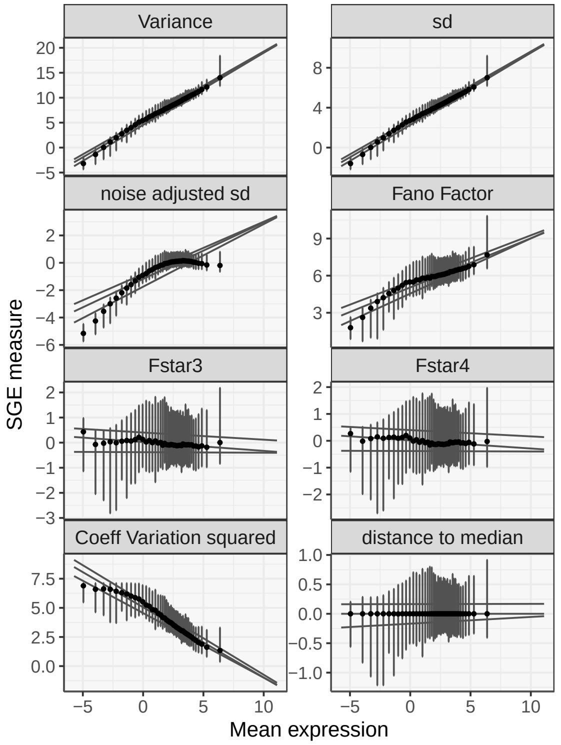


**(b) *Mus musculus* (Liver)**

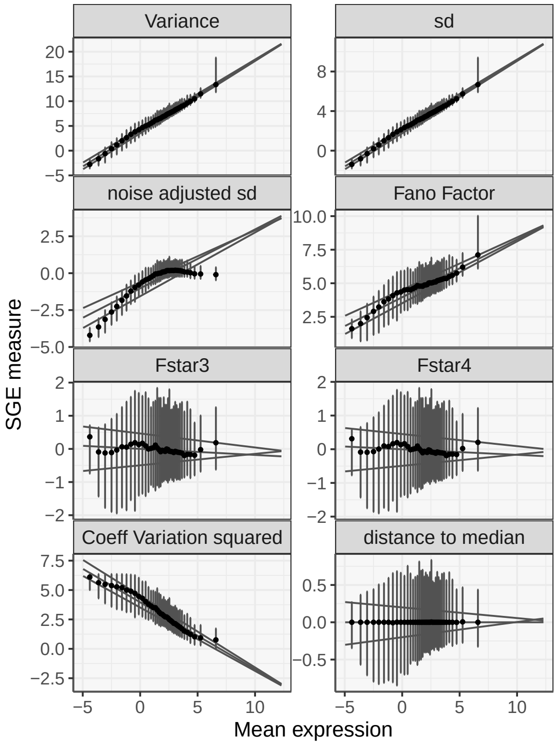


**(d) *Mus musculus* (Aorta)**

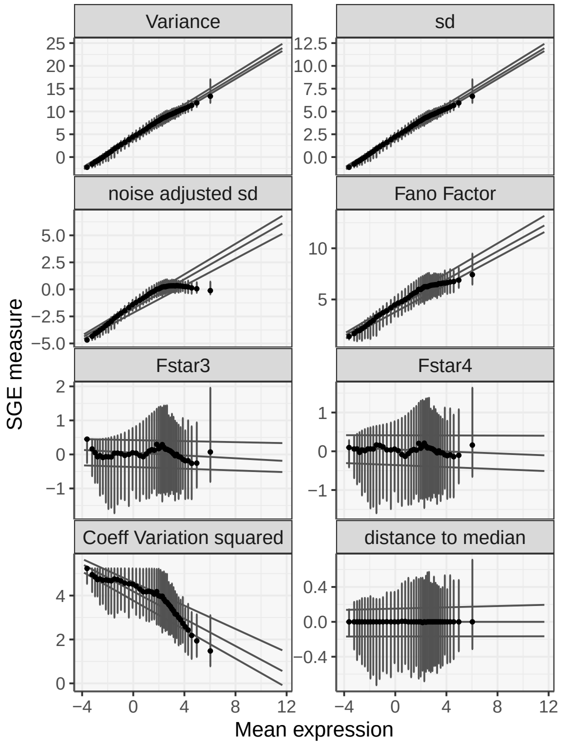

**(e) *Mus musculus* (Heart)**

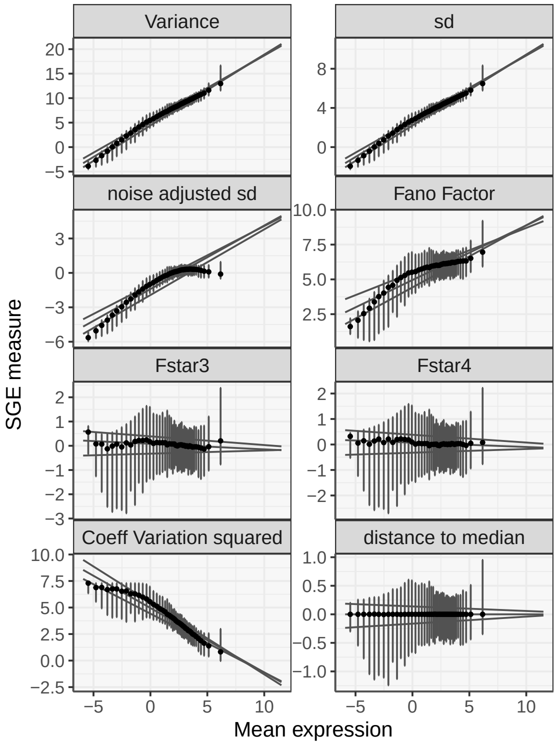


**(g) *Mus musculus* (Muscle)**

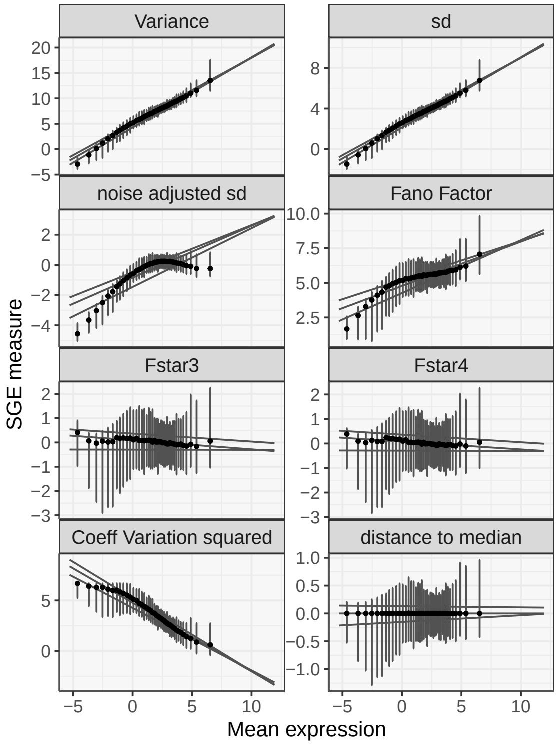


**(f) *Mus musculus* (Kidney)**

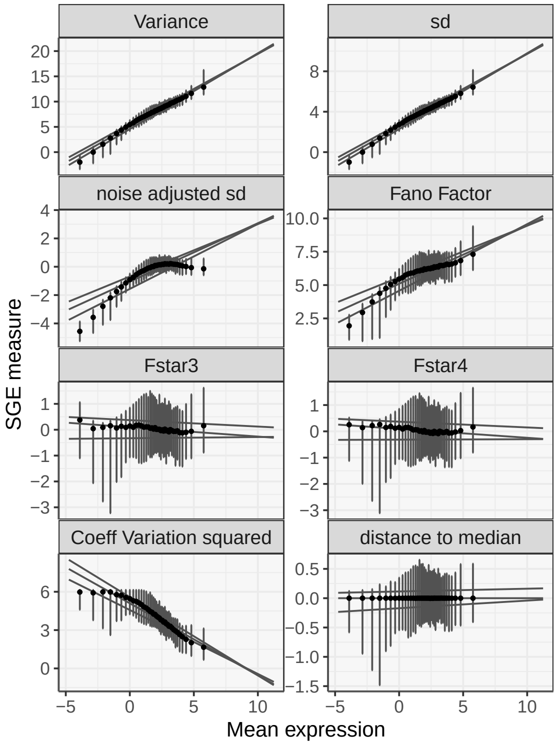


Figure **S20**: **a-f)** Relationships between different stochastic gene expression (SGE) estimations and the mean gene expression, using discretive display with error-bars that represent the middle 95% of values. The 3 horizontal lines separate the values in 4 quartiles. The dots represent the median of the values.
All parameters are log transformed. $F*$ is the noise estimated by the method of Barroso et al. (Barroso et al. 2018). $F*$_min is the first polynomial degree which break the correlation between the noise with mean expression, and $F*$_max is the next one.
