## Supplementary figures for "Two levels of selection of rhythmicity in gene expression: energy saving for rhythmic proteins and noise optimization for rhythmic transcripts"

S1-S7

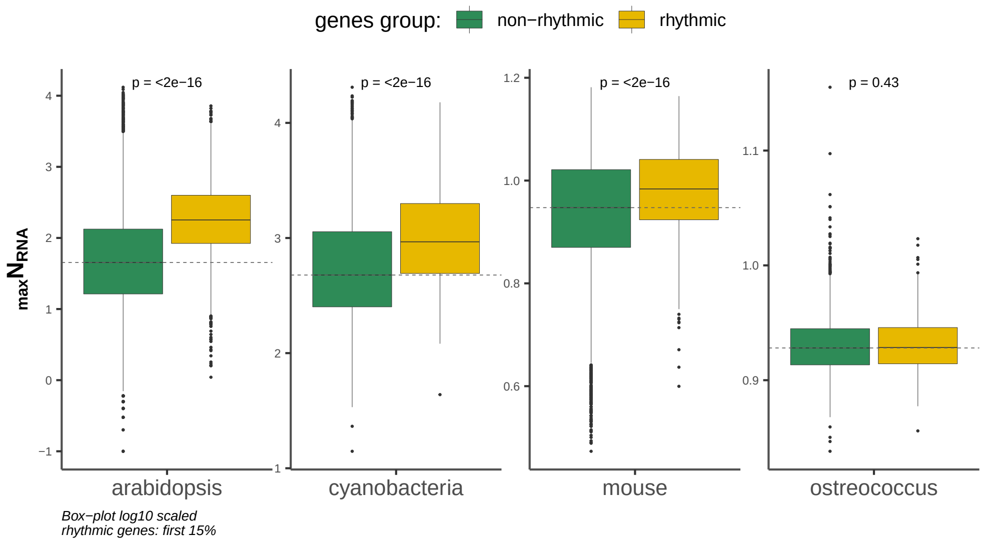


(a) Maximum RNA level over time-points (See Methods)

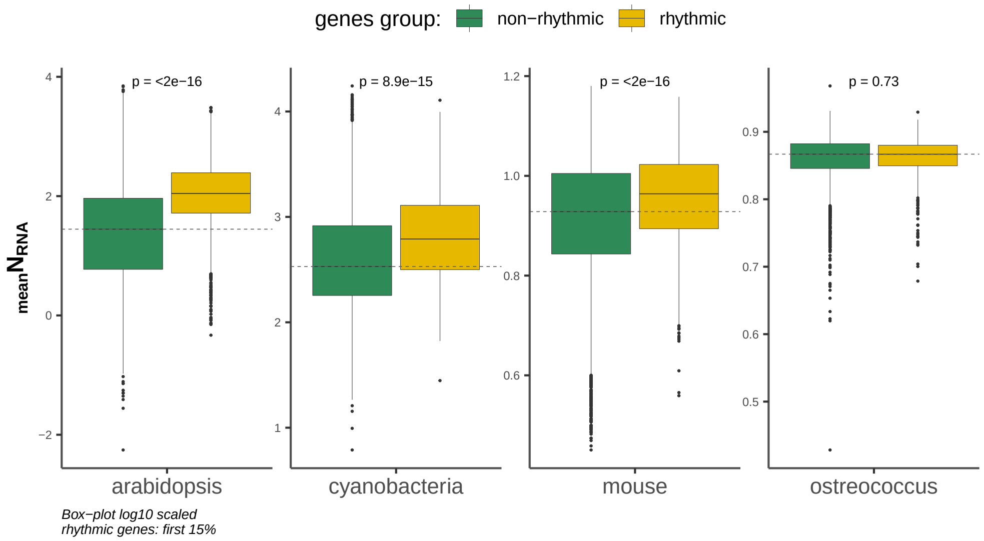


(b) Mean RNA level over time-points

Figure **S1**: Mean or maximum mRNA expression level calculated from time-series datasets. Rhythmic transcripts are highly expressed transcripts.

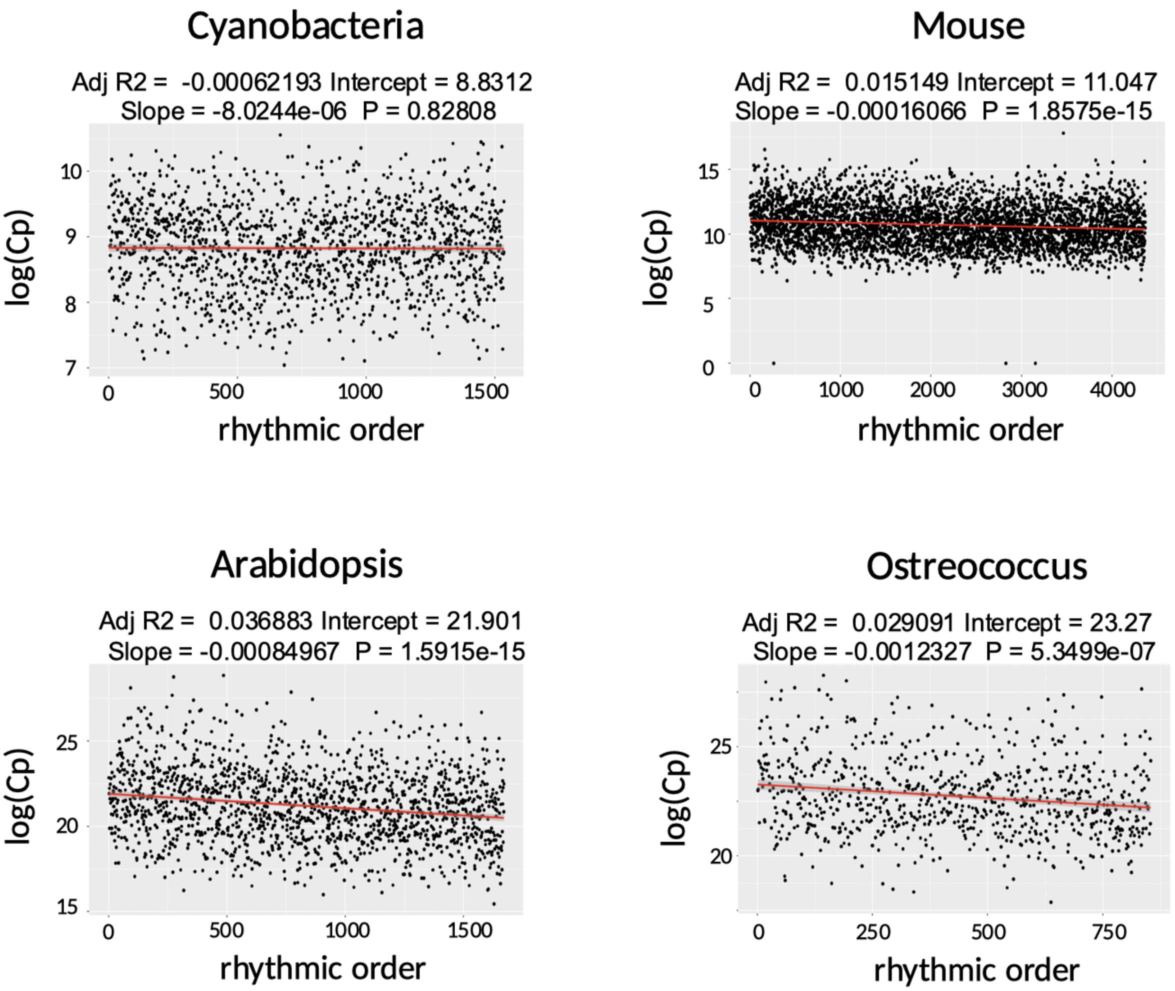


Figure **S2**: Linear regression analysis between the order of the rhythmicity signal (from the most rhythmic to the most un-rhythmic genes) and the total expression cost ($C_p$, calculated from the mean expression level).

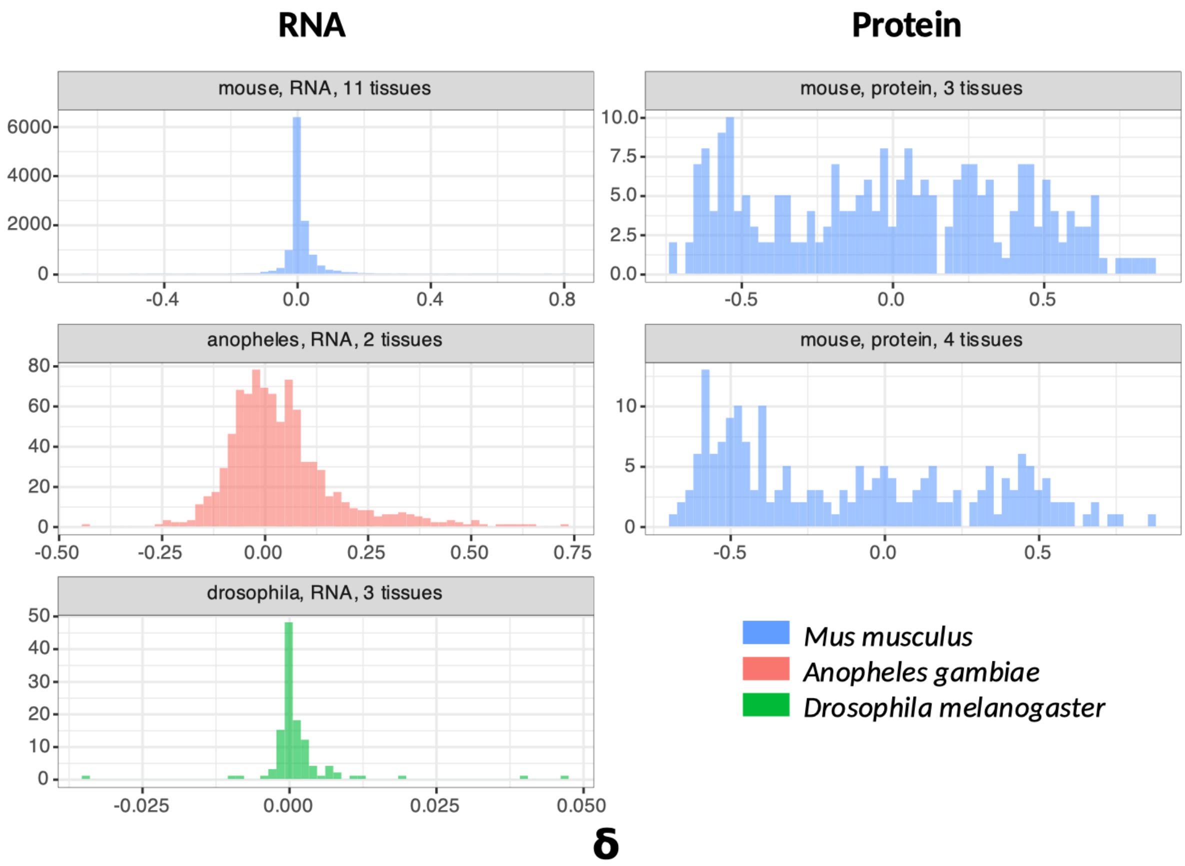


Figure **S3**: Distribution of the difference of expression levels ($\delta$) between rhythmic tissues group and non-rhythmic tissues group obtained for every gene ($\delta$ calculated from mean expression levels; see Methods). $\delta$ shows a bimodal distribution at protein level.

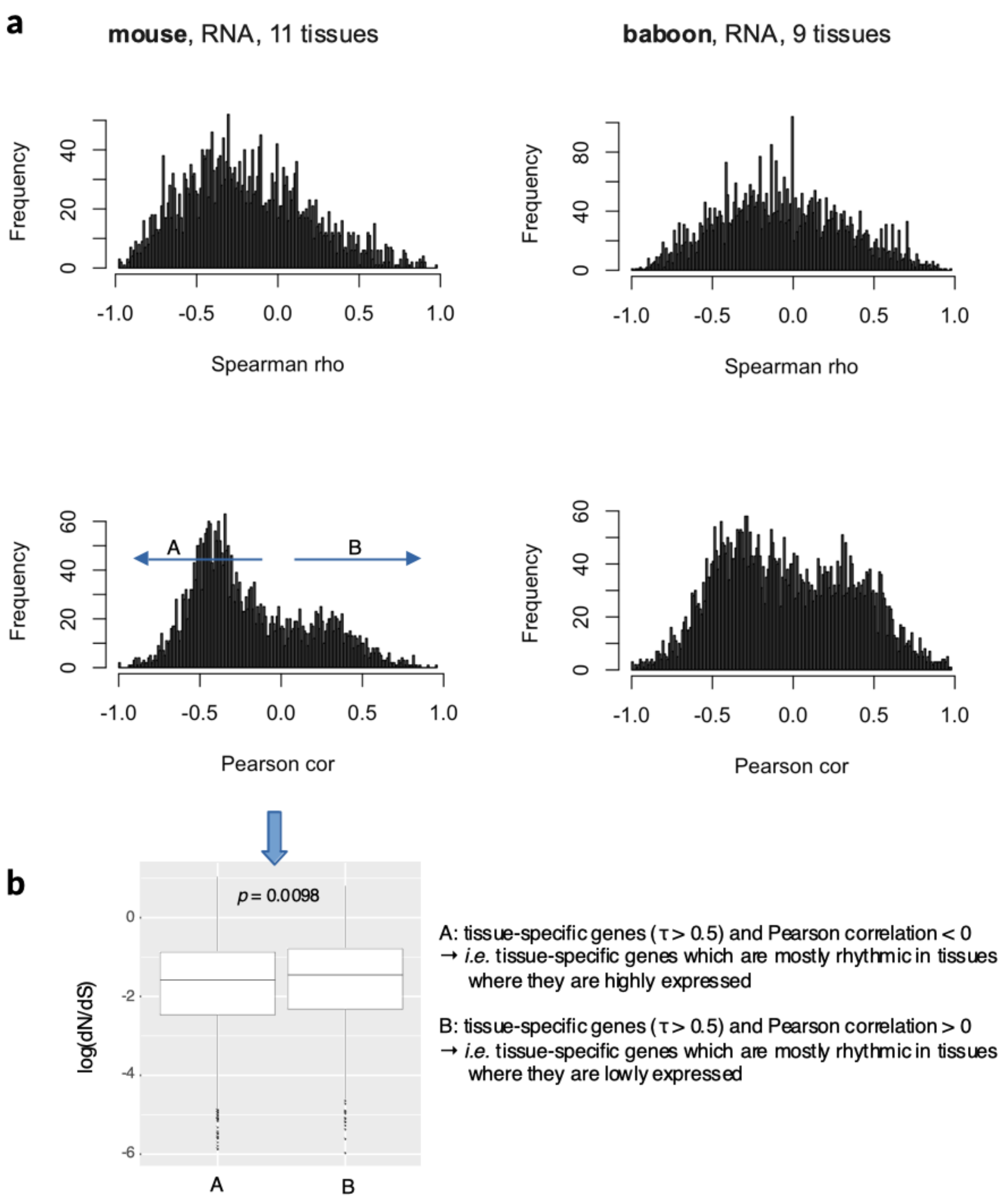


Figure **S4**: **a)** Histograms of Pearson's and Spearman's coefficients testing the correlation between the expression level and the rhythmicicity signal (rhythm $p$-value) measured across the body for every tissue-specific genes ($\tau$ >0.5). **b)** Student's t-test comparing the mean of dN/dS between gene sets A and B: Tissue-specific genes which are mostly rhythmic in tissues where they are highly expressed are under stronger selective constraint than those which are rhythmic in tissues where they are lowly expressed.

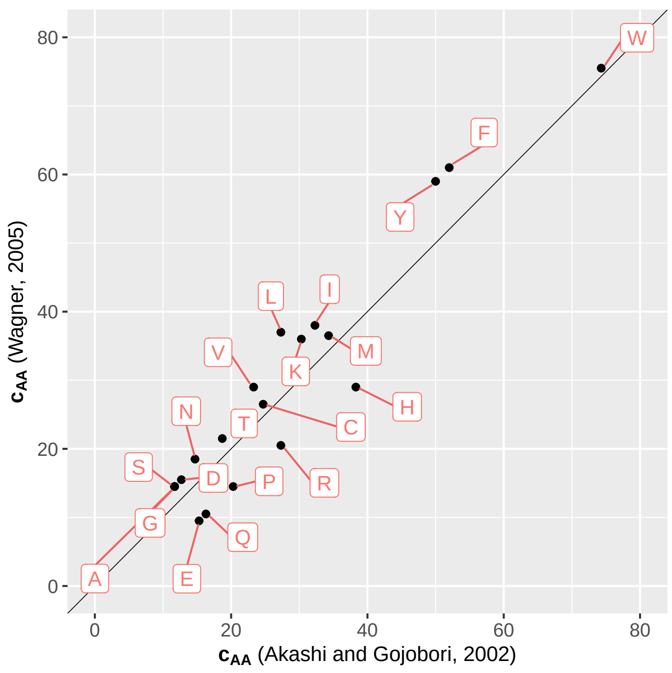


Figure **S5**: Linear relationships of amino acid (AA) biosynthesis costs estimated by Akashi and Gojobori [1], and Wagner [2].

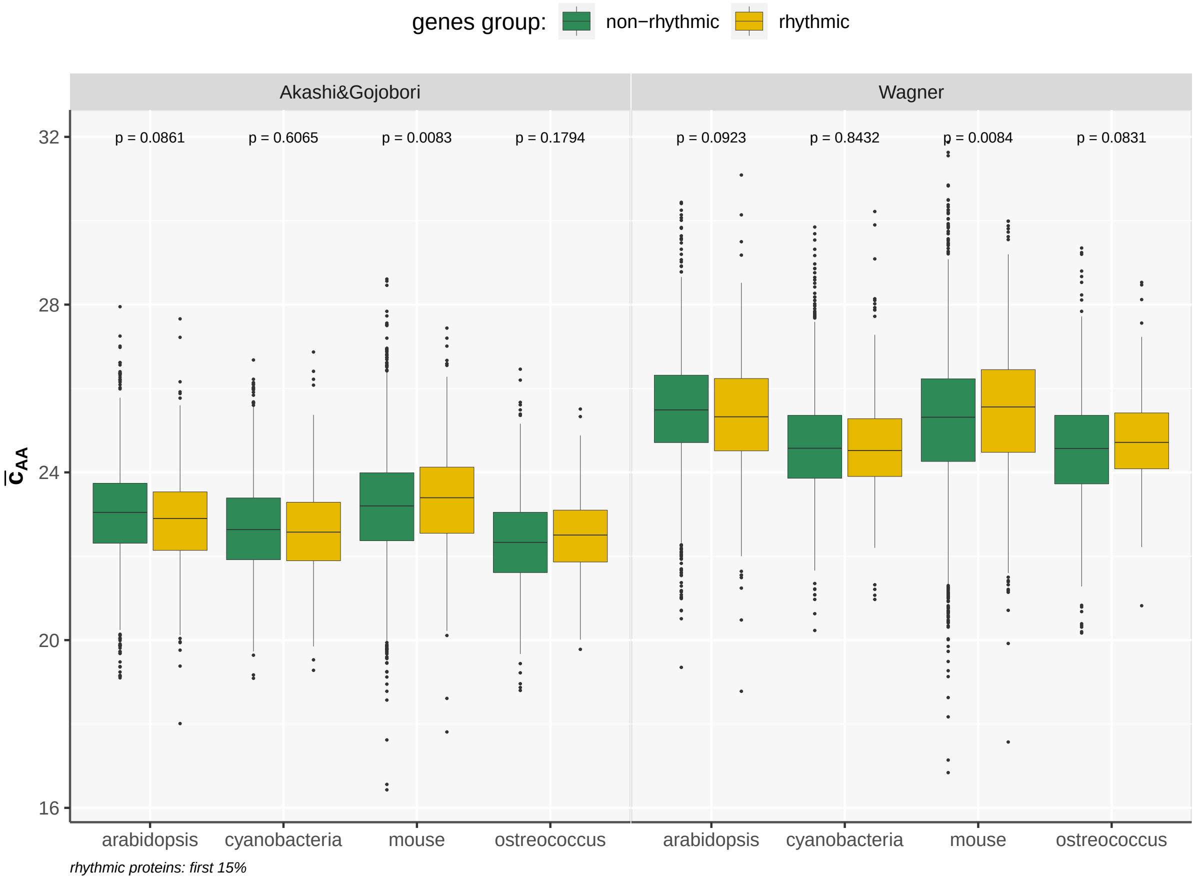


Figure **S6**: Comparison of the averaged AA synthesis costs calculated in both groups: rhythmic VS random genes group, using Akashi and Gojobori versus Wagner AA costs data.

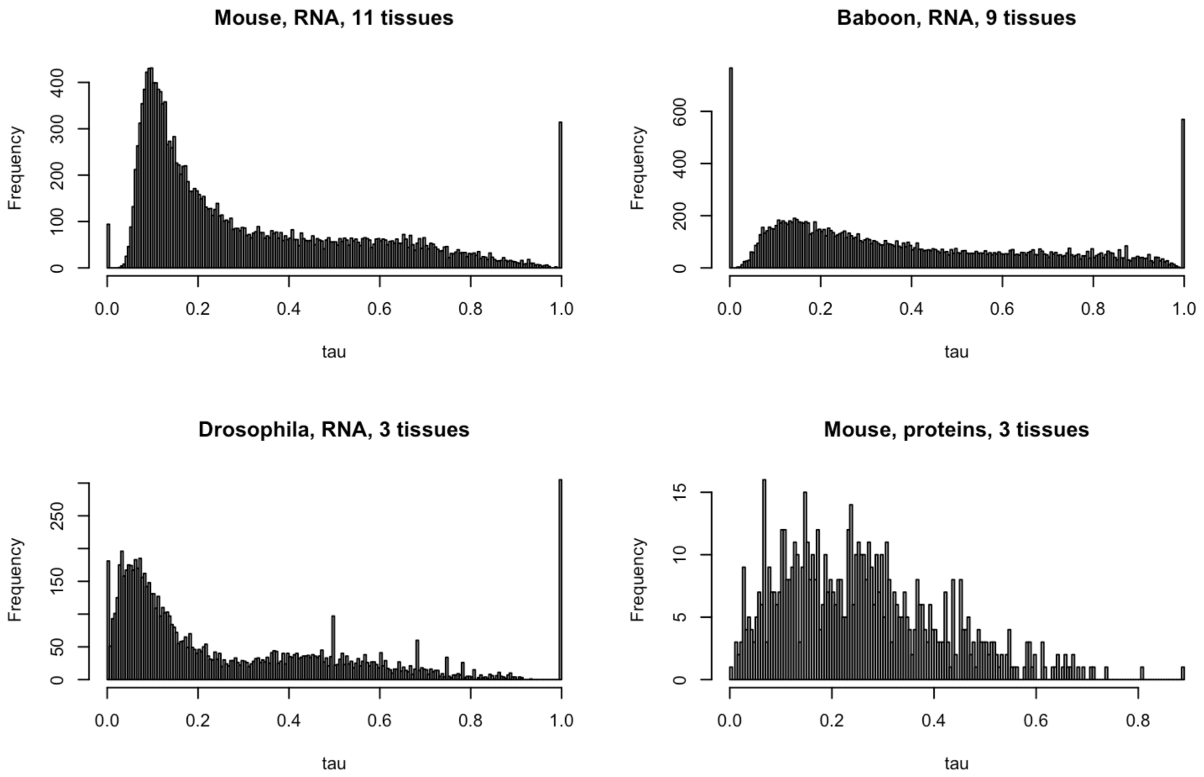


Figure **S7**: Histograms of tissue-specificity $\tau$.
